# Cell autonomous TGF-beta signaling is essential for cell recruitment into degenerating tendons

**DOI:** 10.1101/2020.11.11.378505

**Authors:** Guak-Kim Tan, Brian A. Pryce, Anna Stabio, Douglas R. Keene, Sara F. Tufa, Ronen Schweitzer

## Abstract

Understanding the role of cell recruitment in tendon disorders is critical for improvements in regenerative therapy. We recently reported that targeted disruption of TGFβ type II receptor in the tendon cell lineage (*Tgfbr2^ScxCre^*) resulted in tenocyte dedifferentiation and tendon degradation in post-natal stages. Here we extend the analysis and identify direct recruitment of stem/progenitor cells into the degenerative mutant tendons. Cre-lineage tracing indicates that these cells are not derived from tendon ensheathing tissues or from a *Scleraxis*-lineage, and they turned on tendon markers only upon entering the mutant tendons. Through immunohistochemistry and inducible gene deletion, we further find that the recruited cells originated from a *Sox9*-expressing lineage and their recruitment was dependent on cell-autonomous TGFβ signaling. These results thus differ from previous reports of cell recruitment into injured tendons, and suggest a critical role for TGFβ signaling and cell recruitment in the etiology and treatment of tendon degeneration.

## Introduction

Response to tissue damage or pathology commonly involves the activation or recruitment of stem/progenitor cells that help to replenish the tissue and, in many cases, participate in the healing response (Rennert et al., 2012). While some stem/progenitor cells reside within the tissue, e.g. satellite cells in muscle (Yin et al., 2013), in other scenarios they are recruited from an external niche (Mathews et al., 2004; Jujo et al., 2010; Xynos et al., 2010). Direct detection and investigation of tissue specific stem/progenitor cells was revolutionized by the advent of Cre technology that facilitated the ability to label specific cell populations and monitor their involvement in healing and pathology (Kan et al., 2018; Harvey et al., 2019). The emerging theme is one where for each tissue there are various types of stem/progenitor cells that participate in such processes and likely reflect differential responses to different types of injury or pathology (Rinkevich et al., 2011; Marecic et al., 2015). Identifying the specific cells that participate in the healing response and the signals involved in their activation and recruitment is critical for progress in efforts to enhance and improve the clinical outcomes of therapy. In this study, we identify a new type of progenitor cell involved in the response to tendon pathology.

Tendons are type I collagen rich tissues that transmit the forces generated by muscle contraction to bone (Kannus, 2000). The considerable mechanical challenges to tendons result in a high frequency of injuries that range from acute damage, e.g. in tendon laceration, to chronic damage due to overuse and tissue degeneration as in tendinopathy (Sharma and Maffulli, 2005; Cook et al., 2016). The considerable burden of tendon injuries to individuals and society is compounded by the slow and frequently poor healing of these tissues that often results in impaired tissue integrity (Gerber et al., 2000; Boileau et al., 2005). A better understanding of biological processes underlying tendon healing may thus provide insight towards more effective therapies for tendon injuries.

Experimental investigation on tendon repair has mainly focused on acute injury using transected animal tendons (Forslund and Aspenberg, 2003; Ferry et al., 2007; Howell et al., 2017). Just like other tissues within the body, it has been suggested that cells involved in tendon healing can be from both intrinsic and extrinsic sources (Biro and Bihari-Varga, 1974; Harrison et al., 2003; Jones et al., 2003; Howell et al., 2017). In the latter scenario, recent studies have shown that some of the recruited cells express stem/progenitor markers (Dyment et al., 2014; Runesson et al., 2015; Wang et al., 2017; Harvey et al., 2019). Moreover, several groups have reported that stem/progenitor cells can be isolated from the surrounding peritenon (i.e. epitenon and paratenon) and tendon sheath (Mienaltowski et al., 2013; Wang et al., 2017), and suggest that these tissues may be a source of the recruited cells. Indeed, by taking advantage of *Cre/loxP* reporter system for cell lineage tracing, Dyment *et al* reported that alpha-smooth muscle actin (α-SMA)-positive paratenon cells are the major contributor to the healing response following patellar tendon injury (Dyment et al., 2014). Other lineage tracing experiments have indicated the potential involvement of TPPP3 and osteocalcin-expressing cells during tendon repair (Wang et al., 2017; Harvey et al., 2019). Moreover, injured mouse Achilles tendon was found to be infiltrated by stem/progenitor cells that exhibited different regional distribution and temporal expression (Runesson et al., 2015), implying the existence of multiple recruited cell populations. Despite this recent progress in understanding of the healing response in tendons, basic biology of recruited cells including identity, signals responsible for recruitment, and their role at the injured site, remains largely unknown.

We recently reported that disruption of TGFβ signaling in the tendon cell lineage by targeting TGFβ type II receptor gene *Tgfbr2* using the *ScxCre* driver (*Tgfbr2^ScxCre^*) resulted in tenocyte dedifferentiation in early postnatal stages. Tendon cells in *Tgfbr2^ScxCre^* mutants appeared normal during embryogenesis, but in early postnatal stages lost all differentiation markers including Scx, tenomodulin and collagen I (Tan et al., 2020). Extending the analysis of these mutant tendons, we now find that mutant tendons also began to show progressive degenerative changes, a feature frequently observed in tendinopathy (Kannus and Jozsa, 1991; Longo et al., 2018). Moreover, we find that cells with stem/progenitor features were recruited into the deteriorating mutant tendons. The recruited cells originated from a *Sox9*-expressing lineage, and we further demonstrate that TGFβ signaling was essential for their recruitment. Additionally, it appears that these cells are different from those reported in other studies of cell recruitment into tendons, suggesting a unique cell population that is implicated in cell recruitment into a degenerating tendon.

## Results

### ScxGFP-expressing cells in Tgfbr2^ScxCre^ mutant tendons are newly recruited

Targeting of the TGFβ type II receptor gene *Tgfbr2* with *ScxCre* resulted in a dramatic tendon phenotype (Tan et al., 2020). In early postnatal stages, a few lateral tendons disintegrated and snapped, while in the majority of the tendons, the tenocytes lost the tendon cell fate and dedifferentiated (Figure 1A) (Tan et al., 2020). Macroscopically, the mutant tendons appeared grey and thin, in contrast to normal tendons in which the tight organization of the collagen fibers results in a brilliant white color with firm texture (Figure 1B). The underlying changes were examined by transmission electron microscopy (TEM) and histological analyses that revealed disorganization of collagen matrix, severe disruption of the epitenon structure and paratenon thickening (Figure 1C-G). These tendons may therefore share some similarities with tendons in various pathological conditions (Longo et al., 2008; Dyment et al., 2013).

**Figure 1.**
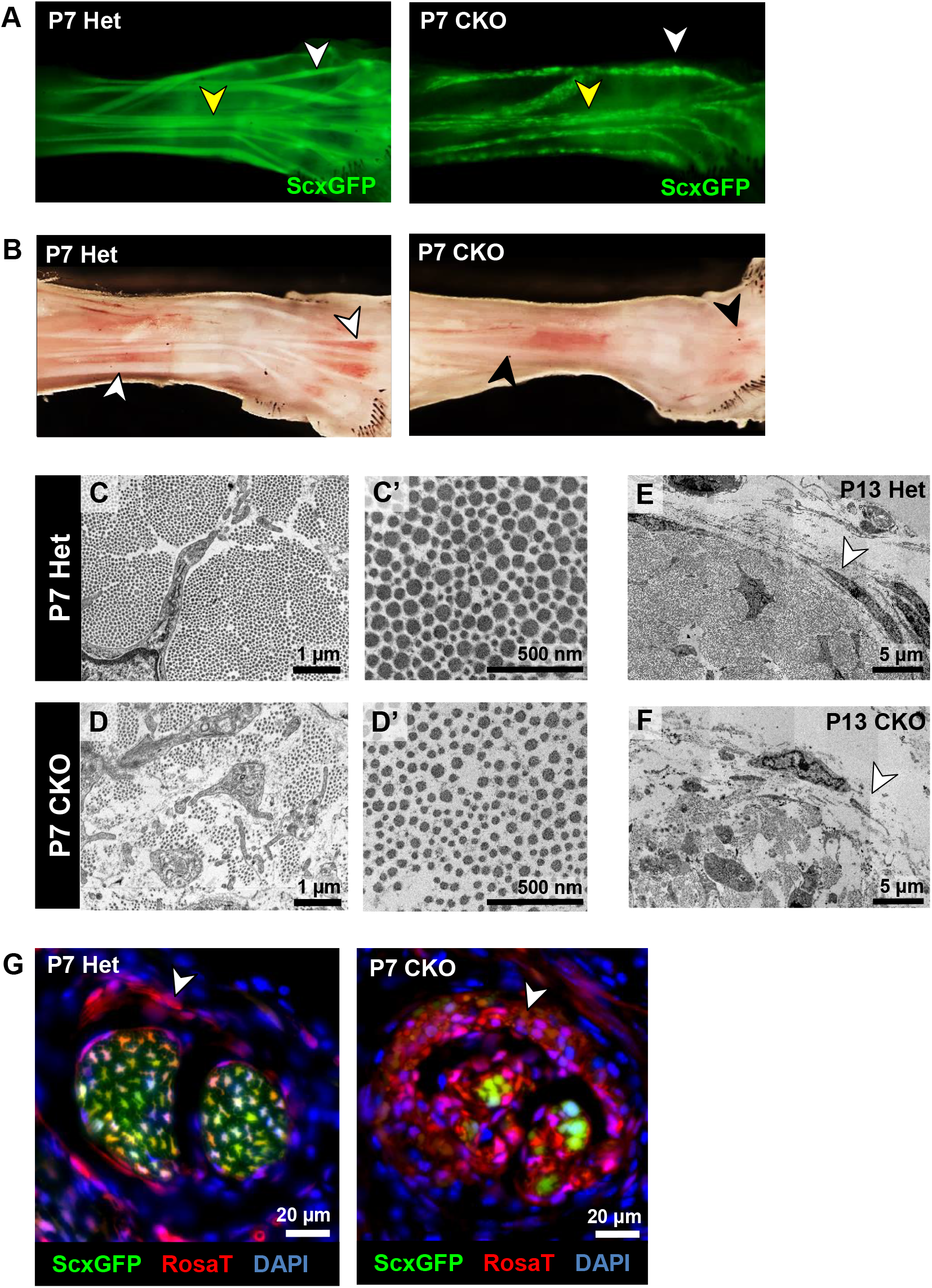
Disruption of tendon structure in *Tgfbr2^ScxCre^* mutant. (A) Comparison of tendon reporter *ScxGFP* signal in forelimbs from P7 heterozygous control (*Tgfbr2^f/+^;ScxCre*) and mutants pups revealed that in mutants a few lateral tendons were missing (white arrowhead) and in the other tendons there was a substantial loss of the *ScxGFP* signal (yellow arrowhead). (B) Brightfield imaging of skinned forelimbs from P7 heterozygous control and mutant pups. While normal tendons display a brilliant white color reflecting the tight organization of the collagen fibers (white arrowheads), mutant tendons had a pale grey appearance (black arrowheads), likely reflecting disruptions to the collagen matrix. (C-F) TEM analysis of heterozygous control and mutant tendons. (C,D) In P7 normal tendons the collagen matrix was highly organized while in mutant tendons the collagen fibrils were smaller with significant gaps between fibril bundles. C’ and D’ are high-magnification images of collagen fibrils in C and D, respectively. (E-F) By P13, in some regions of mutant tendons the epitenon was disrupted and discontinuous (white arrowheads). (G) Triple labeling of transverse sections of heterozygous control and mutant tendons using DAPI nuclear counterstain, *ScxGFP* and Cre reporter *Ai14 Rosa26-tdTomato* (*RosaT*). *RosaT* labeling of the paratenon highlights significant thickening of the tissue observed in some mutant tendons (white arrowheads). Mutant: CKO, Heterozygous: Het.

While the majority of cells in the degenerating tendons of *Tgfbr2^ScxCre^* mutants lost expression of *ScxGFP* and other tendon differentiation markers by P7 (Figure 2A, black arrowhead) (Tan et al., 2020), we observed the appearance of cells that contrary to the surrounding cells expressed *ScxGFP* (Figure 2A, white arrowhead). The *ScxGFP*-positive cells in mutant tendons also expressed other prototypic tendon markers tenomodulin and *Col1a1* (Figure 2E,F, white arrowheads). Despite the induction of tendon markers, these cells differ morphologically from normal tenocytes at this stage. P7 wild-type tenocytes display a stellar-like morphology and a rectangular shape in transverse and longitudinal sections, respectively (Figure 2B,D, black arrowheads). In contrast, the *ScxGFP*-positive cells appeared large and rounded in both views (Figure 2A,C, white arrowheads). Factors that contribute to the aberrant morphology of these cells were not identified to date.

**Figure 2.**
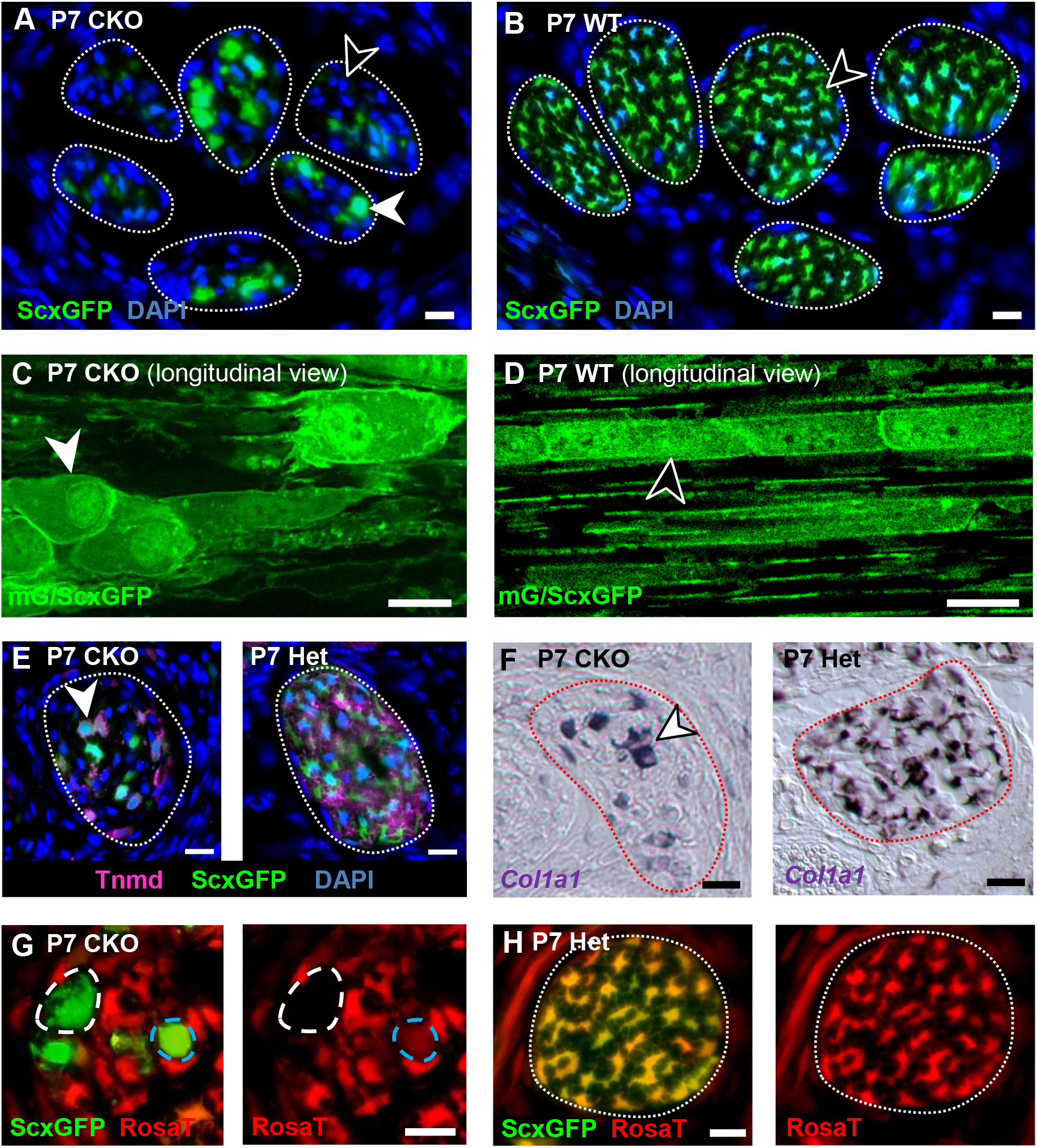
Morphology and marker expression of the *ScxGFP-positive* cells in *Tgfbr2^ScxCre^* mutant tendons. Transverse sections (A,B,E-H) and longitudinal sections (C,D) of tendons from forelimbs of P7 wild-type and mutant pups. In all panels, *ScxGFP* identifies cells with tendon gene expression and nuclear DAPI staining reflects general cellular distribution. (A,B) In wild-type pups all cells of the extensor digitorium communis (EDC) tendons expressed the *ScxGFP* reporter. Conversely, the majority of cells in mutant tendons lost *ScxGFP* expression (black arrowhead), but a small number of *ScxGFP-*expressing cells were also found in these tendons (white arrowhead). Interestingly, while the wild-type tendon cells had a star-like morphology in transverse section, the *ScxGFP*-positive mutant tendon cells were significantly larger and had a round morphology. (C,D) Longitudinal sections from EDC tendons of P7 wild-type and mutant pups carrying the *ScxGFP* and *ScxCre;mTmG* reporters. While *ScxGFP* labels the cell body of tenocytes, *ScxCre;mTmG* results in membrane *GFP* signal to further accentuate the cell morphology (Muzumdar et al., 2007). Wild-type tenocytes were organized in prototypic cell rows and had a rectangular shape in longitudinal view (black arrowhead). However, the *ScxGFP-* positive cells in mutant tendons were rounded and the row organization was disrupted (white arrowhead). (E,F) The *ScxGFP*-positive cells in mutant tendons also expressed tendon markers tenomodulin and *Col1a1*. (E) Immunofluorescence for tenomodulin (Tnmd) in P7 forelimb tendons. Note that in heterozygous control tendons all tenocytes expressed Tnmd but in mutant tendons most cells lost Tnmd expression and only the *ScxGFP*-positive cells expressed Tnmd. (F) *In situ* hybridization for *Col1a1* on tendons from P7 mutant and heterozygous pups. While all tendon cells expressed *Col1a1* in heterozygous control, only a handful of large and rounded cells were positive in mutant tendons (white arrowhead). (G,H) *ScxGFP* and *ScxCre;RosaT;ScxGFP* expression in tendons from mutant and heterozygous control pups. (G) Some of the *ScxGFP*-positive cells exhibited weak or no expression of the Cre reporter *RosaT* (blue and white circles respectively), suggesting these cells may be newly recruited with a recent induction of *Scx*. (H) Lower magnification image of representative P7 control tendons showing all tendon cells were marked by robust *RosaT* expression at this stage. Dashed lines demarcate tendons. Scale bar, 10 μm. Mutant: CKO, Wild-type: WT, Heterozygous: Het.

Surprisingly, some of these *ScxGFP*-positive cells exhibited weak or no expression of the Cre reporter *Ai14 Rosa26-tdTomato* (*RosaT*) (Figure 2G). Conversely, all tendon cells in P7 *Tgfbr2^f/+^;ScxCre* heterozygous pups were marked by robust *RosaT* expression (Figure 2H). *ScxGFP* and *ScxCre* are transgenic mice that utilize the *Scleraxis* enhancer to drive expression of *eGFP* and *Cre* respectively (Pryce et al., 2007; Blitz et al., 2013). In mice that carry both constructs, e.g. the *Tgfbr2^ScxCre^* mutant, it is likely that *ScxGFP* signal will be detected first upon *Scx* activation, since detection of *RosaT* requires an intermediate step of protein synthesis. First Cre activity has to reach threshold levels to induce reporter recombination followed by a second step in which the reporter signal is accumulated to achieve detectable levels. Based on this logic, we hypothesized that *ScxGFP*-positive but *RosaT*-negative cells in mutant tendons (called hereafter *ScxGFP+;RosaT−*) are cells from non *Scx*-expressing cell lineage in which the *Scx* enhancer was recently activated, i.e. they were newly recruited into the mutant tendons. Indeed, when *ScxGFP+;RosaT−* cells were isolated by fluorescence activated cell sorting (FACS) and cultured, they subsequently also showed expression of the *RosaT* reporter (Supplementary Figure 1).

Direct detection of cell recruitment in the process of tendon healing, as demonstrated by this observation, is exciting because it may open new directions for analysis of the healing response. We therefore wanted to reinforce this result with an approach that will identify newly recruited cells by a positive signal rather than the absence of expression of a reporter. To achieve this goal we repeated the experiment utilizing the *mTmG* dual fluorescent Cre reporter in which the ubiquitously expressed membrane-tomato (*mT*) is replaced by membrane-GFP (*mG*) upon Cre-mediated recombination (Figure 3A) (Muzumdar et al., 2007). The advantage of this reporter system over *RosaT* is that it allows a simultaneous visualization and determination of both the recombined and non-recombined states. As expected, in P7 tendons, *mTmG* cells were labelled red (*mT*) in the absence of Cre activity (Figure 3B), while all tendon cells were recombined and appeared positive for *mG* in *ScxCre;mTmG* pups (Figure 3C). On the other hand, in tendons of the *Tgfbr2^ScxCre^* mutant some of the *ScxGFP*-positive cells had a recombined Cre reporter (Figure 3D, white arrowhead), whereas others retained the *mT* signal indicating they did not recombine the reporter or at least have not yet lost the *mT* signal (Figure 3D, yellow arrowhead).

**Figure 3.**
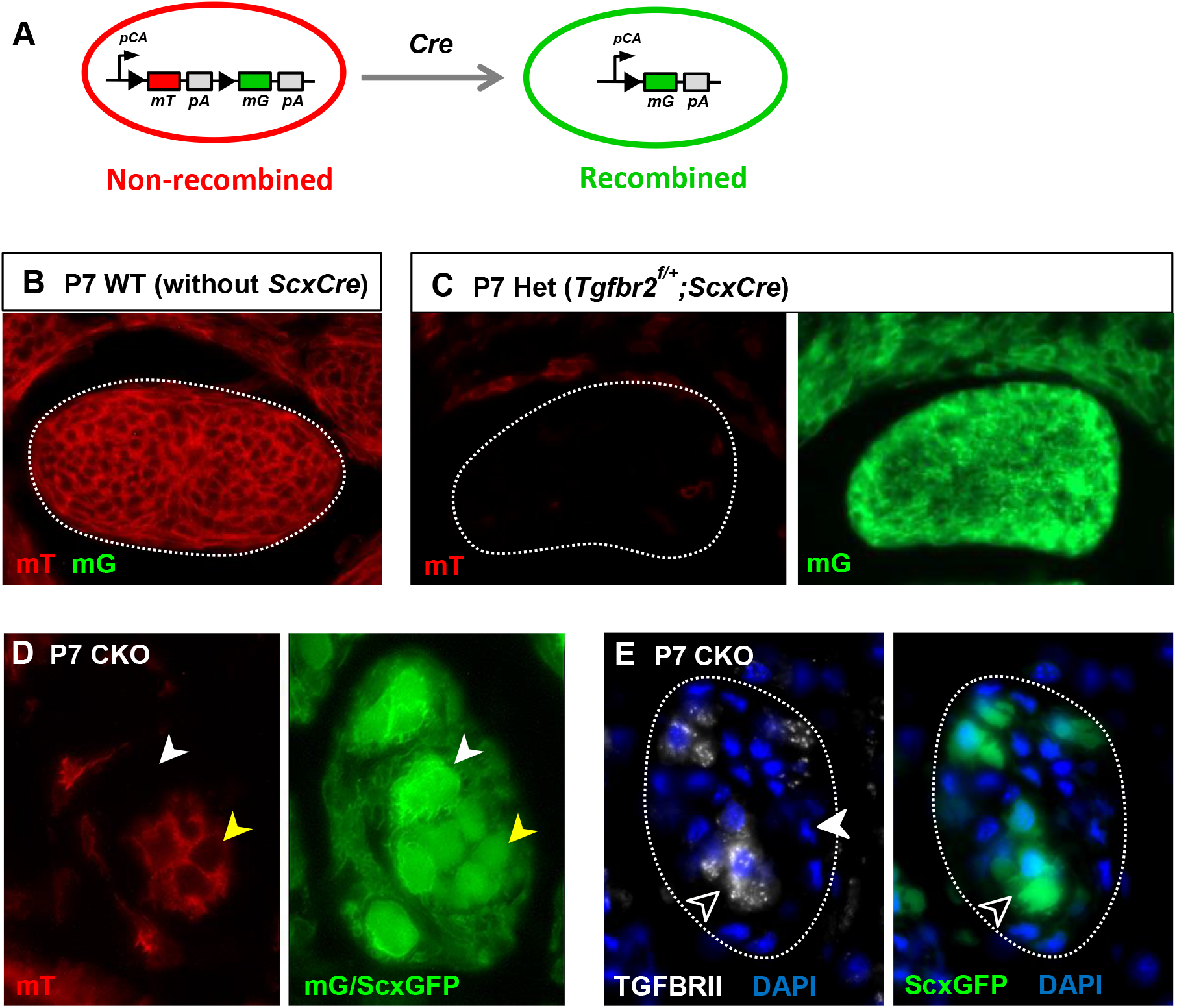
*ScxGFP*-expressing cells in *Tgfbr2^ScxCre^* mutant tendons are newly recruited. (A) Schematic illustration of the *mTmG* dual-fluorescent Cre reporter (Muzumdar et al., 2007). Ubiquitously expressed membrane-tdTomato (*mT*) is replaced by membrane-GFP (*mG*) upon Cre-mediated recombination, allowing simultaneous detection of the recombined and the non-recombined states. (B-E) Transverse sections of P7 forelimb tendons. (B,C) *mTmG* expression in wild-type tendons. In the absence of Cre activity, red (*mT*) fluorescence signal is detected in all cells including tendon cells. (C) In *Tgfbr2^f/+^;ScxCre;mTmG* heterozygous pups, tendon cells were recombined and switched the fluorescence from red to green (*mG*). (D) In *Tgfbr2^ScxCre^* mutant tendons, the *mTmG* Cre reporter was recombined in some of the *ScxGFP-positive* cells (white arrowheads) whereas the Cre reporter was not recombined in other ScxGFP-positive cells (yellow arrowhead). (E) Immunofluorescence staining with antibodies to the TGFβ type II receptor (TGFBRII) in mutant tendons. The resident tendon cells have completely lost expression of the receptor as expected at this stage (white arrowhead), but some of the *ScxGFP-*positive cells still expressed the TGFBRII, further reinforcing the absence of Cre activity in these cells. Tendons are demarcated by dashed lines. Mutant: CKO, Heterozygous: Het. Images not to scale.

To further evaluate Cre activity in the *ScxGFP*-positive cells we attempted to detect expression of the TGFβ type II receptor protein and indeed found expression of the receptor in some of these cells (Figure 3E, black arrowhead). Taken together, these observations reflect a recent induction of *Scx* expression in the *ScxGFP*-positive cells, suggesting these cells are newly recruited into the mutant tendons.

### Temporal dynamics of cell recruitment into mutant tendons

There are only a handful of reports of cell recruitment into tendons (Dyment et al., 2013; Wang et al., 2017; Harvey et al., 2019; Kaji et al., 2020), and almost nothing is known about the origin of such cells or the mechanisms of their recruitment. A robust method for detecting such cells as *ScxGFP+;RosaT−* cells in the tendons of *Tgfbr2^ScxCre^* mutants therefore provided a unique opportunity to learn more about this process. In tendons of wild-type pups with the same marker combination, nearly all cells were positive for both *ScxGFP* and *RosaT* while *ScxGFP+;RosaT−* cells were very rare (Figure 4A), suggesting this marker combination indeed provides a robust approach for identifying newly recruited cells.

**Figure 4.**
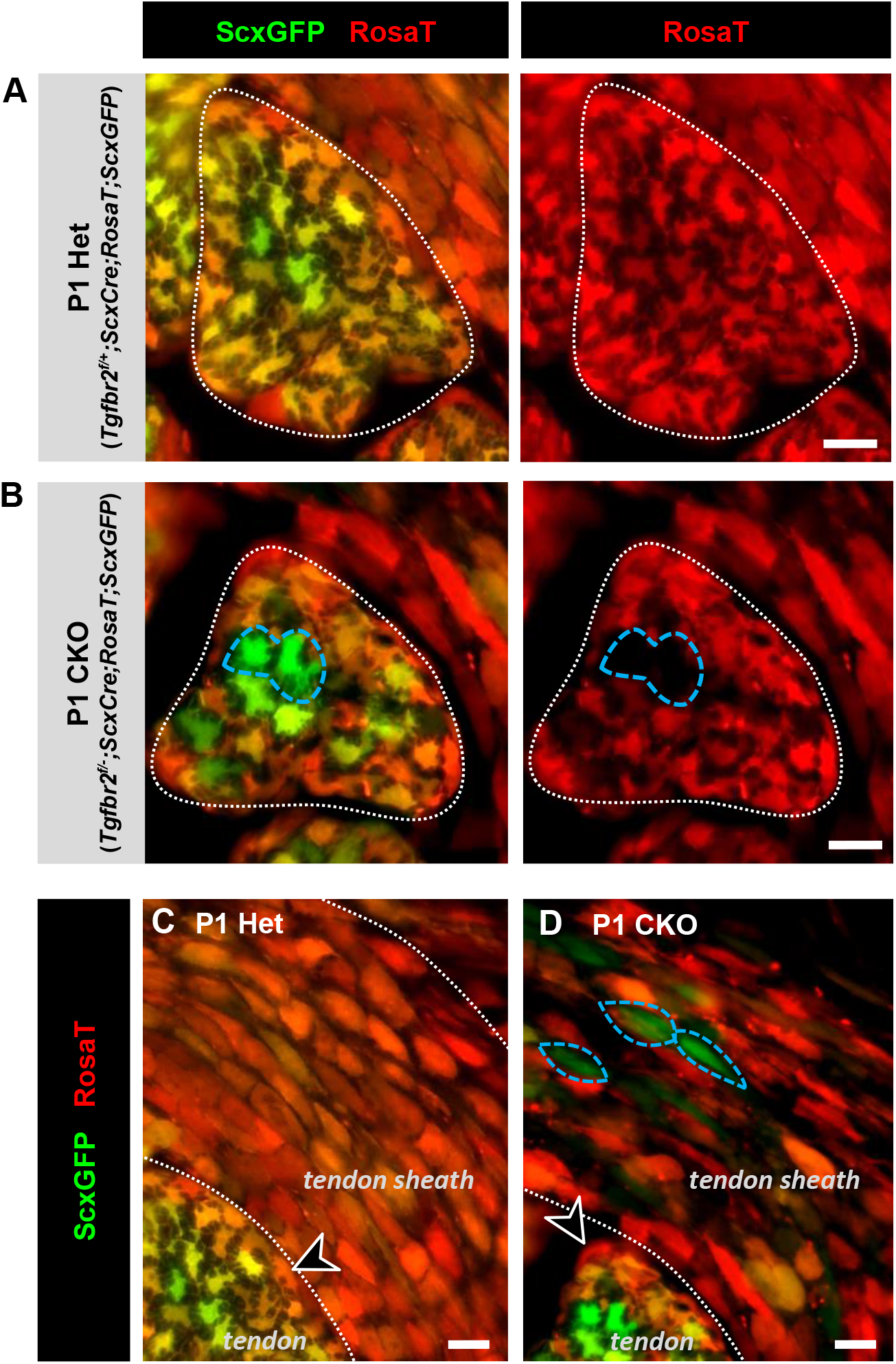
The recruited cells in *Tgfbr2^ScxCre^* mutant tendons do not originate from peritenon or the tendon sheath. (A-D) Transverse sections of extensor digitorium communis tendons from *ScxGFP* and *RosaT*-carrying mutant and heterozygous control pups at P1. (A,B) In tendons of P1 heterozygous mice nearly all cells were positive for both *ScxGFP* and *RosaT* while *ScxGFP+;RosaT−* cells were very rare (A). Conversely, there was a noticeable presence of *ScxGFP+;RosaT−* cells in mutant tendons starting at P1, suggesting these were recruited cells in the mutant tendons. (C) In *Tgfbr2^f/+^;ScxCre* heterozygous pups, *ScxCre* drives recombination of the *RosaT* reporter in all the cells of the epitenon (black arrowhead) and tendon sheath, indicating that cells from these regions are from *Scx*-expressing cell lineage. The absence of *RosaT* expression in cell newly-recruited into the mutant tendons indicates that they are not derived from these regions. (D) In *Tgfbr2^ScxCre^* mutant pups, the newly recruited *ScxGFP+;RosaT−*cells could also be detected in the epitenon and tendon sheath of mutant pups (blue dashed circles). Scale bar, 10 μm. Mutant: CKO, Heterozygous: Het.

Utilizing this approach, we found that the newly recruited cells (i.e. *ScxGFP+;RosaT−*) could be detected already at P1 in the tendons of mutant pups (Figure 4B, blue circle). The levels of these cells within mutant tendons peaked between P1 to P3 and remained detectable throughout the observation period (results not shown). Significantly, in these early postnatal stages most mutant tendons were intact and did not show structural indications of damage (Tan et al., 2020).

Moreover, since tissue repair involves early recruitment of immune and inflammatory cells to the damaged site (Millar et al., 2010; Kragsnaes et al., 2014), we wanted to determine if the recruited cells were associated with an immune response and thus examined for the presence of relevant markers. In both P1 and P7 mutant tendons, we found only a small number of cells expressing the activated macrophage marker F4/80 (Figure 5A). Notably, there was no noticeable difference in their numbers compared to normal tendons (Figure 5B). Moreover, mutant tendons stained negatively for the inflammatory marker TNF-α (Figure 5C), as also observed in normal tendons (Figure 5D). Cell recruitment into the tendons of *Tgfbr2^ScxCre^* mutants thus initiated prior to any sign of a structural destruction or immune response, suggesting a specific molecular signal and not general tissue damage may be the driver of cell recruitment in this case.

**Figure 5.**
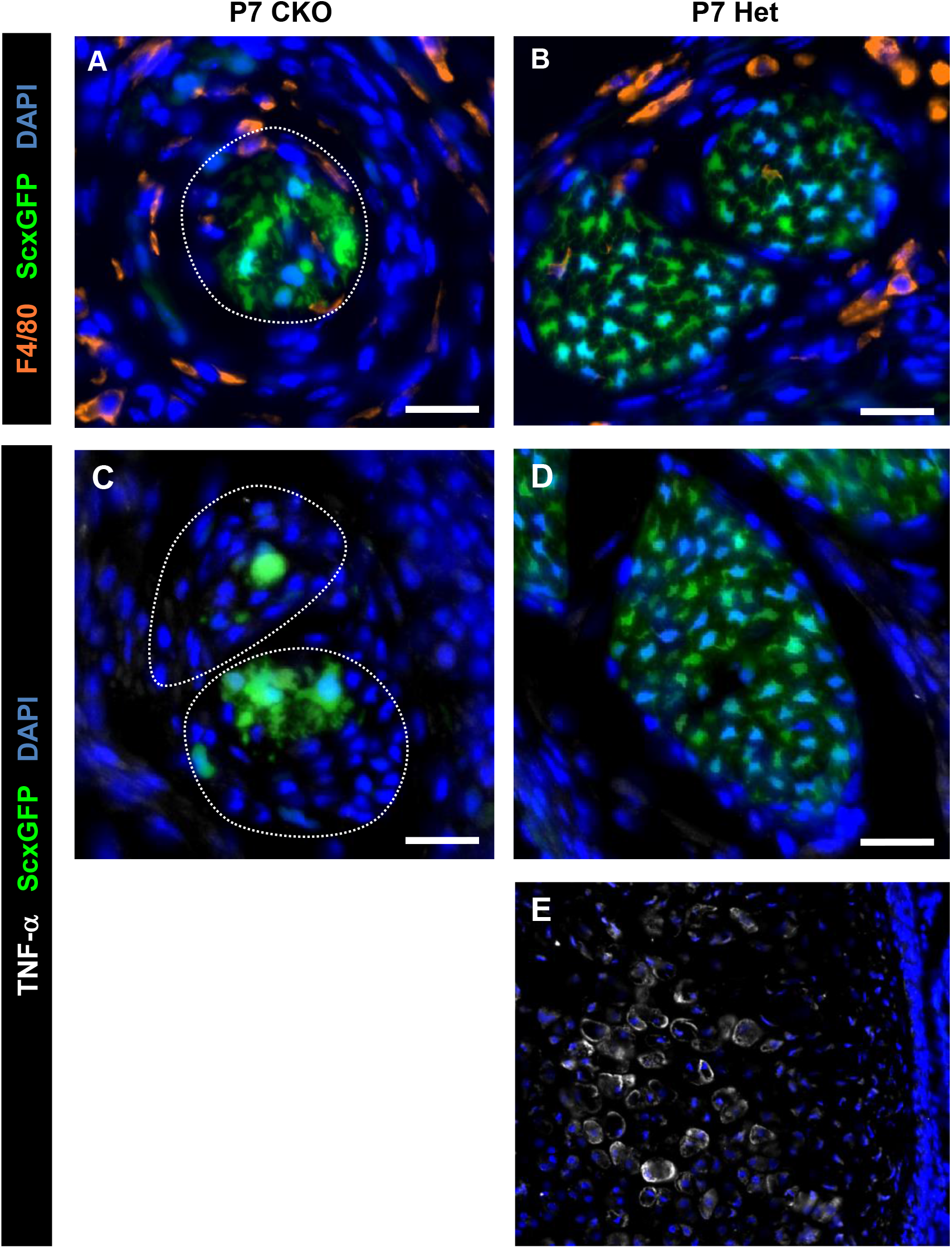
The inflammatory response is not activated in neonatal *Tgfbr2^ScxCre^* mutant tendons. (A-E) Transverse sections of P7 forelimb tendons from mutant and littermate Het controls. In all panels DAPI nuclear staining was used to highlight cell distribution and *ScxGFP* labels tenocytes. (A,B) Immunofluorescence staining for the macrophage-specific antigen F4/80. Only a small number of cells expressed the F4/80 antigen in both the mutant and Het control tendons. (C-E) Immunofluorescence staining for the inflammatory marker TNF-α. (C,D) Expression of TNF-α was not detected in tendons of both the mutant and heterozygous littermate control. (E) Bone marrow region on the same section as in (D) that serves as a positive control for TNF-α staining (image not to scale). Mutant tendons are demarcated by dashed lines. Scale bar, 20 μm. Mutant: CKO, Heterozygous: Het.

### The recruited cells do not originate from peritenon or the tendon sheath

A handful of recent studies identified cell recruitment into tendons mostly in the context of injury (Dyment et al., 2013; Tan et al., 2013; Runesson et al., 2015; Wang et al., 2017) and possibly also following physiological loading (Mendias et al., 2012). While the origin of such cells remains unclear, it was suggested in a few studies that they may arise from the peritenon (i.e. paratenon and epitenon) or tendon sheath (Dyment et al., 2013; Wang et al., 2017; Harvey et al., 2019). To assess if the recruited cells in *Tgfbr2^ScxCre^* mutant tendons were derived from peritenon or tendon sheath, we again took advantage of the *RosaT* Cre reporter system. Not much is known about gene expression in these tendon ensheathing tissues that clearly do not overlap with gene expression in tenocytes (Harvey et al., 2019; Tan et al., 2020). Interestingly, while peritenon and tendon sheath cells do not express the tendon reporter *ScxGFP* they are consistently positive for *RosaT* Cre reporter in mice carrying *ScxCre;RosaT* alleles (Figure 4C). This combination of markers likely represents transient expression of *Scx* in early progenitor cells of the peritenon and tendon sheath that was sufficient for activation of the Cre reporter. Since the newly recruited cells are from non *Scx*-expressing lineage and do not express the Cre reporter *RosaT* as noted earlier (Figure 4B), the cells recruited into the mutant tendons were not derived from peritenon and tendon sheath.

Notably, the newly recruited *ScxGFP+;RosaT−* cells could also be detected in the peritenon and tendon sheath of *Tgfbr2^ScxCre^* mutant pups (Figure 4D, blue circles). Since the absence of Cre reporter expression indicates that these are not original peritenon or tendon sheath cells, we postulate that these are either the recruited cells entering the mutant tendons by passing through peritenon and tendon sheath, or cells also being recruited into these tendon ensheathing tissues in mutant pups.

Additionally, previous studies have demonstrated the invasion of cells expressing α-SMA, also a pericyte marker, into injured tendons with a likely endothelial-perivascular origin (Dyment et al., 2013; Howell et al., 2017). To determine if this may also be the origin of the recruited cells in the tendons of *Tgfbr2^ScxCre^* mutants we examined expression of endothelial and pericyte-associated markers (Cathery et al., 2018), but could not detect expression of CD31, CD146 or α-SMA in these cells (data not shown). Taken together these results suggest that the cells getting recruited into the tendons of *Tgfbr2^ScxCre^* mutants are different from the cells so far reported to be implicated in tendon injury, and possibly represent a repair response to the pathological (degenerating) changes in the *Tgfbr2^ScxCre^* mutant tendons.

### The recruited cells have clonogenic features and express stem/progenitor markers

Direct detection and the ability to isolate the newly recruited cells into the mutant tendons presented a unique opportunity to characterize the cellular features and possibly origin of the recruited cells. We hypothesized that for participation in tendon healing the recruited cells likely have features of stem/progenitor cells and tested for such features. Wild-type tendons contain 2-4% cells with colony forming potential, also known as tendon-derived stem/progenitor cells (TSPC) (Bi et al., 2007; Mienaltowski et al., 2013) (Figure 6B). To test the colony forming capacity of the recruited cells we dissociated cells from the tendons of P7 mutant pups and isolated the recruited cells by FACS based on the unique marker combination of these cells (*ScxGFP*+;*RosaT*-). The cells were seeded at one cell per well in 96-well plates and colony forming potential was determined after 9 to 14 days of culture (Figure 6A). We indeed found that 2.9 ± 0.7 % of the recruited cells had colony forming potential (n = 4 independent experiments in duplicate) (Figure 6B), but the size of clones formed by the recruited cells varied and in general was smaller than the clones of wild-type TSPC (Figure 6C). The difference presumably reflects the dynamic change of cellular state in the recruited cells, in which some cells were newly recruited and still possessed progenitor stemness features, while others have already advanced in assuming the tendon cell fate and thus lost their proliferative capacity to give rise to large colonies.

**Figure 6.**
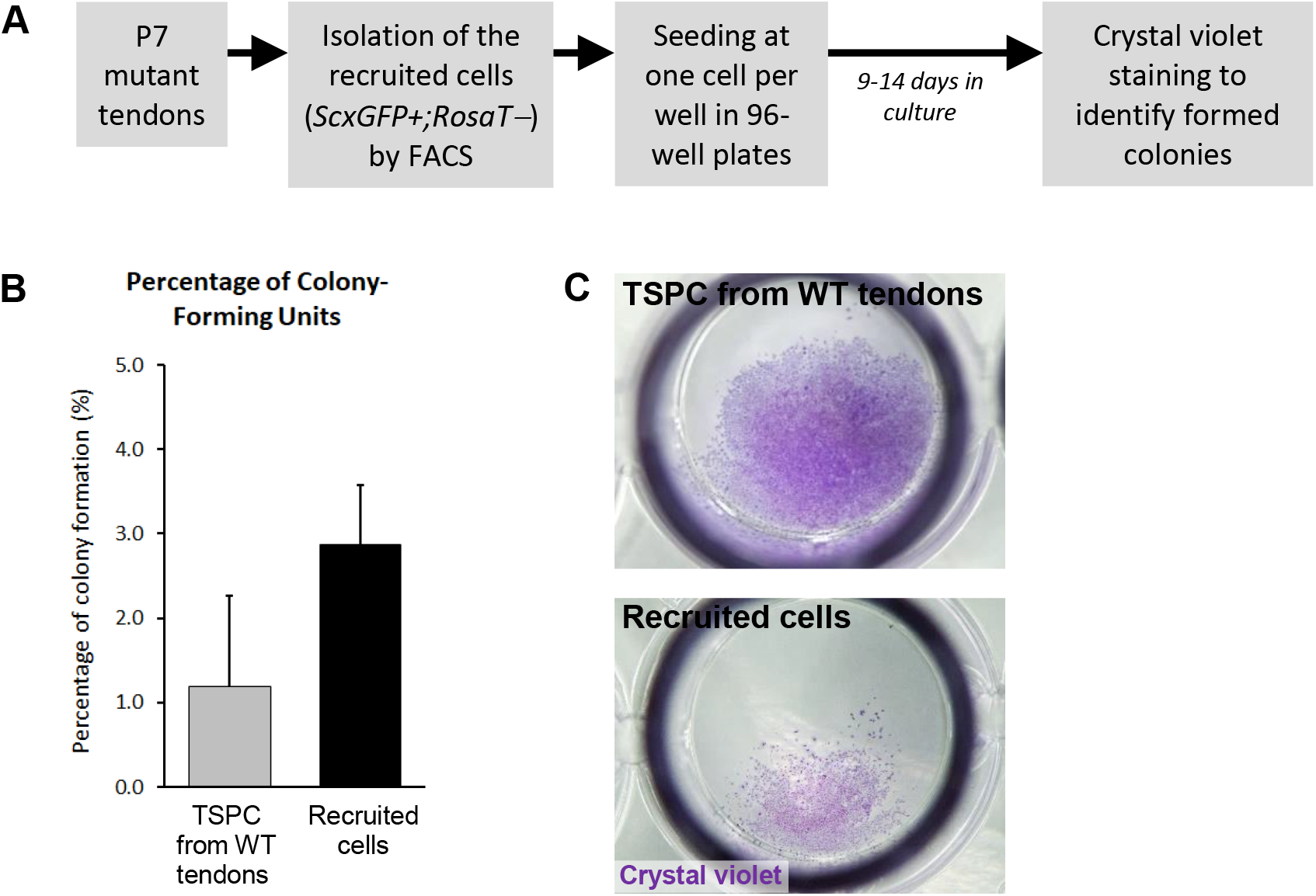
The recruited cells exhibit clonogenic capability in culture. (A) Experimental outline for testing the colony forming capacity of the recruited cells. Briefly, cells were enzymatically-released from the tendons of P7 mutant pups and the *ScxGFP+;RosaT−* newly recruited cells were isolated by FACS sorting. The cells were then seeded at one cell per well in 96-well plates and cultured for 9-14 days. Colony formation was visualized with crystal violet staining and the percentage of wells with colonies served as an indication for colony forming capacity of these cells (Tan et al., 2020). (B) About 2.9 ± 0.7 % of the *ScxGFP+;RosaT−* recruited cells formed colonies in cultures. For a positive control (TSPC), cells were also harvested from wild-type tendons and evaluated for their clonogenic capability under the same culture conditions. The results shown are mean ± SD (n=4 independent experiments in duplicate; a total of 768 cells were analyzed). (C) The size of colonies formed by the recruited cells isolated from *Tgfbr2^ScxCre^* mutants varied and in general was smaller than those of TSPC, presumably reflecting the dynamic change in the stemness state of the recruited cells. Mutant: CKO, Wild-type: WT, Tendon-derived stem/progenitor cells: TSPC.

Recognizing the progenitor state of the recruited cells, we next tested these cells in P7 mutant tendons for expression of typical markers identified in cultured TSPC and other established progenitor markers (Bi et al., 2007; Mienaltowski et al., 2013; Lui, 2015). We first found that the recruited cells expressed the stem/progenitor marker nucleostemin, which is not expressed by normal tendon cells (Figure 7A,C) (Zhang and Wang, 2010). Surprisingly, we also detected expression of Sox9 protein in the recruited cells (Figure 7D). *Sox9* is most commonly recognized as an early cartilage marker but it is also expressed in various populations of stem/progenitor cells (Poche et al., 2008; Scott et al., 2010; Furuyama et al., 2011). It was previously demonstrated that some tendon progenitors express *Sox9*, and some *Sox9CreERT2* activity can be detected in tenocytes even in postnatal stages (Soeda et al., 2010; Blitz et al., 2013; Huang et al., 2019). However, expression of Sox9 protein is undetectable or negligible by immunohistochemistry in normal tenocytes (Figure 7B) and is therefore unique to these recruited cells. Notably, the number of Sox9- and nucleostemin-positive cells was high at P1, coinciding with the earliest detectable recruitment in mutant tendons (Figure 4B) and the prevalence of these cells gradually declined in later stages. Taken together, our results suggest that cells with stem/progenitor features were recruited into mutant tendons. Additionally, the recruited cells expressing the *ScxGFP* reporter were not detected away from the tendons but only within or adjacent to mutant tendons (Supplementary Figure 2), suggesting that irrespective of their origin the cells turned on tendon gene expression only upon entering the tissue.

**Figure 7.**
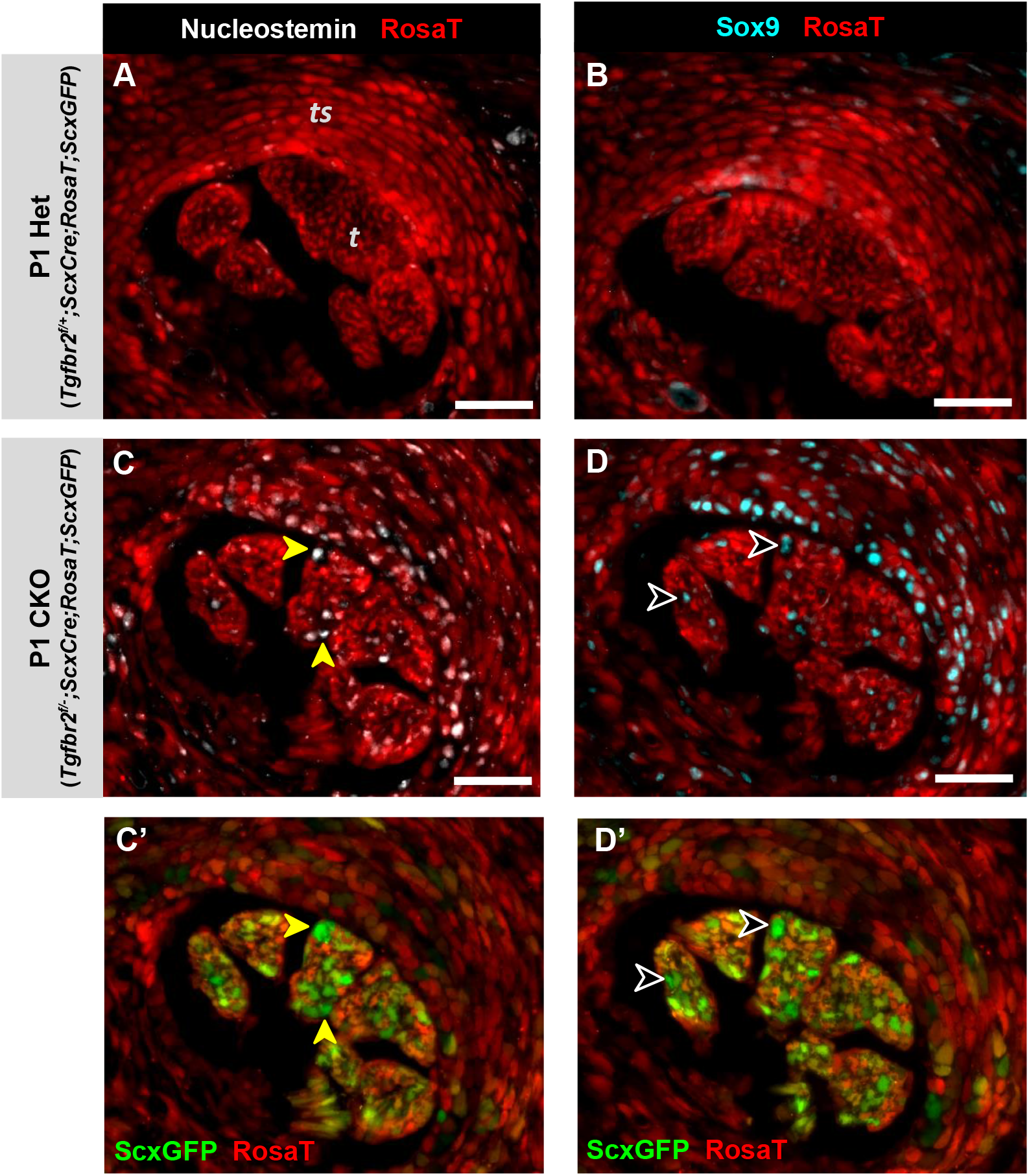
The recruited cells express stem/progenitor markers. Immunofluorescence staining for stem/progenitor markers on transverse sections of extensor digitorium communis tendons from forelimbs of *Tgfbr2^ScxCre^* mutants and heterozygous littermates at P1, a stage with extensive cell recruitment. The *ScxCre* lineage was labeled with the Cre reporter *RosaT*. In heterozygous controls, expression of (A) nucleostemin and (B) Sox9 was either undetectable or negligible in tenocytes and the tendon sheath. (C,D) Conversely, numerous cells expressing both nucleostemin (yellow arrowheads) and Sox9 (black arrowheads) were detected within these regions in *Tgfbr2^ScxCre^* mutants. (C’,D’) represent images from the same field of view in (C,D) respectively, and *ScxGFP* signal was captured to identify the *ScxGFP+;RosaT−* newly recruited cells. Scale bars, 50 μm. Mutant: CKO, Heterozygous: Het, Tendon sheath, ts, Tendons, t.

### Cell autonomous TGFß signaling is essential for cell recruitment into mutant tendons

TGFβ signaling has been implicated in cell motility and recruitment in other systems (Franitza et al., 2002; Tang et al., 2009) and recently also in tendons (Kaji et al. 2020). Since we found that the recruited cells still expressed the TGFβ type II receptor, we next wanted to ask if deletion of *Tgfbr2* in these cells will change their capacity for recruitment. We previously found that the tendon phenotype in *Tgfbr2^ScxCre^* mutants is dependent on the specific spatio-temporal features of *ScxCre* activity (Tan et al., 2020). To target the *Tgfbr2* receptor before the cells are recruited, we therefore decided to add the ubiquitous inducible Cre deletor (*RosaCreERT2*) (Hameyer et al., 2007) to the *Tgfbr2^ScxCre^* allele combination. Since the tendon phenotype manifests in *Tgfbr2^ScxCre^* mutants in postnatal stages, we can use *RosaCreERT2* to induce ubiquitous loss of the receptor at that stage and examine the effect on cell recruitment into mutant tendons.

Pups of the mutant allele combination, *Tgfbr2^f/-^;ScxCre;RosaCreERT2* (called hereafter *ScxCre;RosaCreERT* Double Cre mutant), were given tamoxifen at the earliest time-point of detectable recruitment at P1 and P2, and harvested at P7. Interestingly, we found nearly 60% reduction in the number of recruited cells in the *ScxCre;RosaCreERT* Double Cre mutant pups compared to *Tgfbr2^ScxCre^* mutants (Figure 8A-C) (*p*<0.01; n=3). The dramatic reduction in cell recruitment suggests that TGFβ signaling indeed plays a role in cell recruitment. However, the partial reduction of recruitment in these experiments may also imply the existence of alternative molecular mechanisms or may simply reflect partial Cre activation. Tamoxifen application in neonates has severe deleterious effects. It was therefore not possible to increase the dosage or number of days in which tamoxifen was administered. To test if Cre activity was partial in the *ScxCre;RosaCreERT* Double Cre mutants, we stained mutant forelimb sections with anti-TGFβ type II receptor antibody. Intriguingly, cells that were still recruited into the mutant tendons in the Double Cre experiment were positive for the receptor (Figure 8D, arrowheads), suggesting a complete dependence of cell recruitment on TGFβ signaling. Moreover, the fact that *Tgfbr2* expressing cells could still be recruited in this scenario demonstrates that the loss of TGFβ signaling did not have a general effect on the capacity of cells to be recruited or on the recruiting signal, but rather TGFβ signaling acts cell-autonomously and was required for the ability of individual cells to be recruited in this scenario.

**Figure 8.**
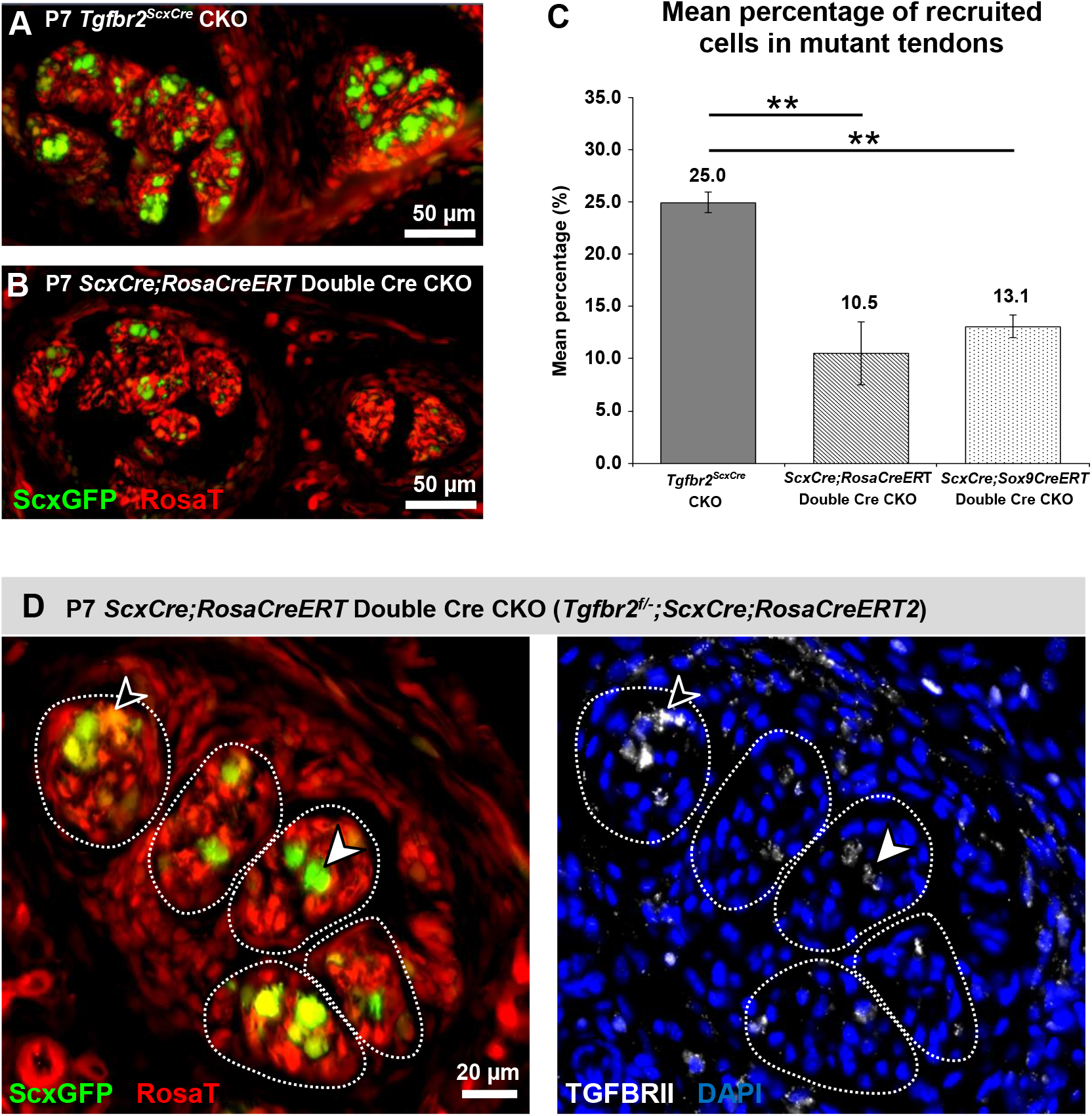
Cell autonomous TGFβ signaling is essential for cell recruitment into mutant tendons. The ubiquitous *RosaCreERT2* driver was incorporated into the *Tgfbr2^ScxCre^* mutant background (*Tgfbr2^f/-^;ScxCre;RosaCreERT2*, called hereafter *ScxCre;RosaCreERT*) to examine the effect of global loss of the TGFβ signaling on cell recruitment into *Tgfbr2^ScxCre^* mutant tendons. Tamoxifen was administered at P1 and P2, and the effects on cell recruitment were evaluated at P7. (A,B) Transverse sections through the extensor digitorium communis tendons from single Cre (*Tgfbr2^ScxCre^*) and *ScxCre;RosaCreERT* Double Cre littermates. Qualitative histological analysis reveals a dramatic reduction in the number of recruited cells (*ScxGFP*-expressing cells) in the Double Cre mutant pups compared to *Tgfbr2^ScxCre^* mutants. (C) Quantitative determination of the number of recruited cells in P7 experimental pups. There was a nearly 60% reduction in cell recruitment in the *ScxCre;RosaCreERT* Double Cre mutant pups, suggesting that TGFβ signaling is essential for cell recruitment. For cell lineage study, the Double Cre strategy was also employed to target *Tgfbr2* specifically in *Sox9*-expressing cells (*Tgfbr2^f/-^;ScxCre;Sox9CreERT2*, called hereafter *ScxCre;Sox9CreERT*). The results showed about 48% decrease in recruited cell numbers, suggesting that the recruited cells are from *Sox9*-expressing cell lineage. The results shown are mean ± SD (n=3, \*\**p*<0.01). (D) Immunofluorescence staining for TGFBRII on transverse sections of *ScxCre;RosaCreERT* Double Cre mutant forelimbs. The left panel shows the overlay of the green (*ScxGFP*) and red (*RosaT*) channel, and the right panel shows for the same section the expression of the TGFBRII on the background of DAPI nuclear counterstain. The *ScxGFP*-expressing cells that were still recruited into Double Cre mutant tendons were also positive for the receptor (arrowheads), suggesting a complete dependence of cell recruitment on TGFβ signaling. Mutant: CKO.

The identity and anatomical origin of the recruited cells is of great importance for future efforts to manipulate and enhance the healing processes. The experimental paradigm used above provided us with a unique tool to test hypotheses regarding the origin of the recruited cells in this experimental model, since we can use various other inducible Cre lines with a more restricted target population to target the receptor and test the effects on cell recruitment. We demonstrated above that most of the newly recruited cells expressed the Sox9 protein (Figure 7D). It was important, however, to determine if *Sox9* expression was induced only during the recruitment process or if it was expressed in the cells prior to their recruitment and therefore may be used as a marker to identify the origin of these cells. We therefore employed the same Double Cre strategy to target *Tgfbr2* but in this time specifically in *Sox9*-expressing cells using a *Sox9CreERT2* driver in combination with the *Tgfbr2^ScxCre^* mutant (*Tgfbr2^f/-^;ScxCre;Sox9CreERT2*, called hereafter *ScxCre;Sox9CreERT* Double Cre mutant). The results showed about 48% decrease in recruited cell numbers (*p*<0.01; n=3) (Figure 8C), suggesting that most if not all of the recruited cells indeed expressed *Sox9* prior to their activation.

## Discussion

The present studies extend our previous observations where targeted disruption of TGFβ signaling in tendon cells (i.e. *Scx*-expressing cell lineage) led to loss of their cell fate (Tan et al., 2020), and provide three major observations. First, we find in post-natal stages a progressive tissue degeneration in the mutant tendons, a condition that has been often associated with tendinopathy and spontaneous tendon rupture (Kannus and Jozsa, 1991; Jarvinen et al., 1997; Longo et al., 2008). Secondly, we identify direct recruitment of stem/progenitor cells into the deteriorating tendons. Furthermore, findings from the Cre-lineage tracing indicate that these cells are not derived from surrounding peritenon or tendon sheath, implying the existence of multiple sources for stem/progenitor cell recruitment in tendons. Thirdly, we find that most if not all of the recruited cells were from *Sox9*-expressing lineage, and TGFβ signaling is essential for their recruitment into the degenerating tendons. This scenario thus opens an opportunity to directly examine the origin of recruited stem/progenitor cells and the mechanisms of their activation in degenerative tendon pathologies.

In mutant tendons, cell recruitment was at a pick already at P1, in the absence of observable structural damage or immune response in these tendons, suggesting that the process of tenocyte dedifferentiation in mutant tendons is also accompanied by the secretion of a specific recruitment signal. We therefore suggest the following model for cell recruitment in this scenario (Figure 9): (1) Tenocyte dedifferentiation in neonatal mutants results also in secretion of a stem/progenitor cell recruitment and/or activation signal(s). (2,3) Activation and/or recruitment of the stem/progenitor cells is dependent on activation of TGFβ signaling in these cells in a cell autonomous manner. TGFβ ligands may therefore be the recruitment signals in this case. It may however also be possible that a different signal is employed for cell recruitment and TGFβ signaling plays an essential role in the activation or motility of the cells towards the degenerating tendons. (4) Expression of *ScxGFP* in the recruited cells is observed only in or near the target tendons, suggesting the induction of the tendon cell fate in these cells is not an integral part of the activation or recruitment process, but rather that an additional local signal or interaction with the tendon cells or environment lead to induction of the tendon cell fate in the recruited cells while they integrate into the mutant tendon.

**Figure 9.**
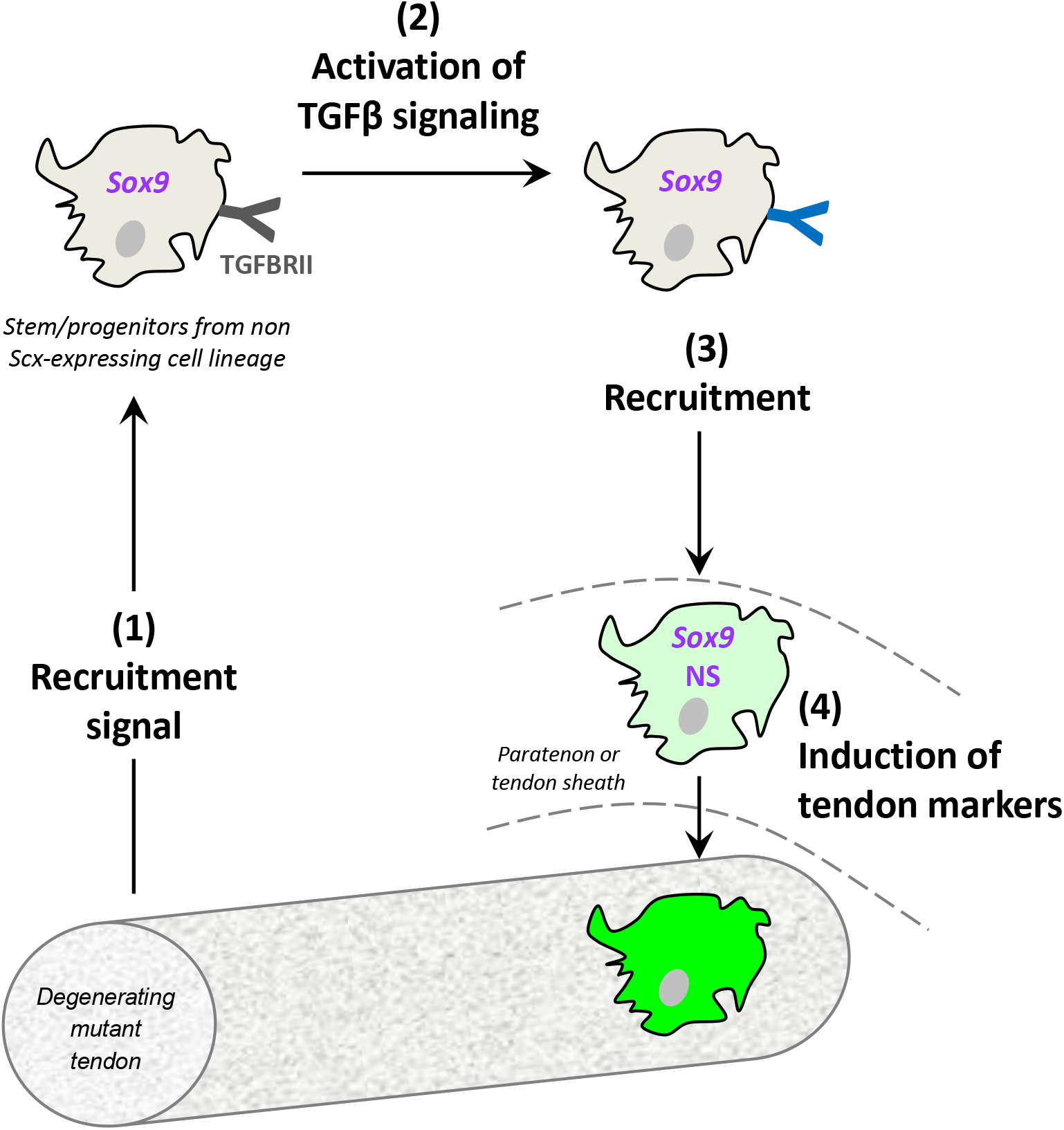
Proposed model for cell recruitment process into *Tgfbr2^ScxCre^* mutant tendons. (1) The degenerating mutant tendons emit a recruitment signal. (2) The recruitment signal leads to activation of TGFβ signaling and to recruitment of a population of *Sox9*-positive and *Scx*-negative stem/progenitor cells. The *ScxCre*-lineage tracing suggests that these cells are not derived from surrounding peritenon or tendon sheath. (3) The activated cells are recruited towards the mutant tendon. (4) These *Sox9*-expressing cells turned on tendon markers and became *ScxGFP*-positive only upon entering the mutant tendons. Notably, the newly recruited cells, identified as *ScxGFP+;RosaT−* cells in this study, could be detected as early as P1 in the mutant tendons, peaked between P1 to P3 and remained detectable throughout the study period. These cells also express nucleostemin (NS), a stem/progenitor marker that has been reported in tendon-derived stem/progenitor cells but not in mature tenocytes. TGFβ type II receptor: TGFBRII.

Tendon damage occurs very frequently, but the tissue tends to heal poorly (Gerber et al., 2000; Boileau et al., 2005). Therefore, there is great interest in the use of stem/progenitor cells to improve tendon repair and therapy (Nourissat et al., 2010; Hernigou et al., 2014; Oh et al., 2014). Previous investigations have demonstrated that acute tendon injury involves recruitment of new cells expressing stem/progenitor markers including α-SMA, Oct-3/4, TPPP3 and osteocalcin (Dyment et al., 2013; Runesson et al., 2015; Howell et al., 2017; Wang et al., 2017; Harvey et al., 2019). However, their relevance to degenerative tendons, a feature that precedes and underlies tendinopathy and tendon rupture, remains unclear. To our knowledge this is the first demonstration of stem/progenitor cell recruitment in the context of a degenerative tendon condition, thus opening new avenues for direct identification and analysis of stem/progenitor cells *in vivo* in the context of tendon pathology.

We show herein that the cells recruited into mutant tendons are clonogenic, and the majority of newly recruited cells are stained positive for nucleostemin and Sox9. Nucleostemin is a GTP-binding protein expressed predominantly in the nucleoli of stem/progenitor cells (Beekman et al., 2006; Nomura et al., 2009). In recent years, nucleostemin has been shown to be expressed by culture-expanded stem/progenitor cells from tendons (Zhang and Wang, 2010). Notably, infiltration of nucleostemin-positive progenitor cells into ruptured rat Achilles tendons has been reported in an earlier study (Runesson et al., 2015). Likewise, *Sox9* is a transcription factor that is associated with various adult progenitor cell populations (Formeister et al., 2009; Furuyama et al., 2011). Lineage tracing on *Sox9CreERT2* mice suggest that *Sox9*-expressing cells also serve as progenitors during tendon development (Soeda et al., 2010; Huang et al., 2019). Importantly, numerous studies have reported the involvement of *Sox9*-positive stem/progenitor cells in tissue repair (Furuyama et al., 2011; He et al., 2017). Our results therefore suggest that *Sox9* is also expressed in the niche of the stem/progenitor cells identified in this study and expression of *Sox9* may therefore serve as an initial indicator for possible location and origins of the cells.

Studies have suggested several possible sources of recruited cells into damaged tendons including peritenon (i.e. paratenon and epitenon) and tendon sheath (Dyment et al., 2013; Wang et al., 2017; Harvey et al., 2019). Apparent changes of epitenon cellular activity, e.g. increased proliferation, has been observed in many cases of tendon injuries (Khan et al., 1996). Moreover, recent studies show appearance and subsequent migration of *ScxGFP*-positive cells from paratenon into tendons following injury (Dyment et al., 2013; Sakabe et al., 2018). A similar phenomenon was observed in this study and prompted us to ask if the recruited cells were derived from these tissues. The results from *ScxCre*-lineage tracing indicate that these cells are derived neither from peritenon or tendon sheath regions. Moreover, the eventual structural destruction of epitenon in our mutant pups may also limit the availability of cells recruited from this region. These results therefore imply the existence of multiple sources of recruited stem/progenitor cells for tendons. Different sources for cell recruitment may reflect specialization of specific cells for different tendon conditions or the concurrent activation of multiple cell populations that may have complementing activities in the healing process. Interestingly, a previous study has shown biphasic infiltration of two different stem/progenitor cell populations into ruptured rat Achilles tendons (Runesson et al., 2015).

To better understand the identity and origin of the recruited cells, further investigation focused on Sox9 because immunostaining demonstrated robust Sox9 expression in these cells during the early phase of recruitment. Results from Double Cre experiment showed that inducible deletion of TGFβ signaling in *Sox9*-expressing cells significantly reduced the number of recruited cells in mutant pups. The finding not only corroborates our earlier notion that the recruited cells expressed Sox9, but further reveals that at least a subpopulation of the recruited cells is from *Sox9*-expressing cell lineage. *Sox9CreERT*-lineage tracing shows the presence of *Sox9*-expressing cells in perichondrium and bone marrow in neonates, suggesting the possibility of these tissues as sources of the recruited stem/progenitor cells. Notably, stem/progenitor cells have been identified in the perichondrium and bone marrow, and these cells are involved in tissue repair (Pineault et al., 2019). The possible involvement in these perichondrial cells in tendon healing will be addressed in future studies.

Studies of stem/progenitor cells and their roles in normal development and pathology were revolutionized with the advent of Cre technology and the ability to label distinct cell populations with a combination of a tissue specific Cre driver and a Cre reporter. These studies are typically prospective, a hypothesis regarding the role of a specific cell population is tested by labeling these cells using a tissue specific Cre (Soeda et al., 2010; Wang et al., 2017; Harvey et al., 2019).

In this study we developed a complementary retrospective approach to identify cell recruitment into tendons. *ScxGFP* is a robust tendon reporter that results in strong GFP expression shortly after activation of the *Scx* enhancer (Pryce et al., 2007). Activation of the *RosaT* reporter in a *ScxCre;RosaT* combination requires two rounds of protein synthesis; First accumulation of sufficient Cre protein and then after recombination and activation of the reporter, accumulation of the reporter protein (Madisen et al., 2010). It is therefore likely that upon induction of the *Scx* enhancer in a cell with no tenogenic history the *ScxGFP* signal will be detected first and the *RosaT* signal will follow with some delay. Notably, this approach will not identify cells from the tendon sheathing tissues or intrinsic tendon cells, since these cells are from *Scx*-expressing cell lineage and thus will have an activated *RosaT* reporter. We suggest however that retrospective screening for *ScxGFP+;RosaT−* cells in tendons of *ScxCre;RosaT;ScxGFP-carrying* mice can be used as a general approach for identification of non *Scx*-expressing cell recruitment into tendons following injury or pathology.

Understanding key players that mediate cell recruitment into pathologic tendons is critical to design tendon reparative strategies. At present almost nothing is known about this process both *in vitro* and *in vivo*. Here we show that TGFβ signaling is essential for the cell recruitment into the degenerating mutant tendon, in which disruption of TGFβ type II receptor in these cells significantly reduced their number in the mutant tendons. TGFβ signaling is known to be involved in the recruitment of various cell types including stem/progenitor cells in pathologic conditions. For instance, blockage of TGFβ signaling with the receptor inhibitor abolished the mobilization and recruitment of mesenchymal stem cells to the injured arteries in mice (Wan et al., 2012). With regard to tendons, activation of TGFβ signaling pathway has been reported during embryonic tendon cell development (Havis et al., 2014). Moreover, a number of studies have demonstrated increased TGFβ ligand and receptor expression by tendon (Fenwick et al., 2001; Dahlgren et al., 2005; Favata et al., 2006) or its adjacent tissues (Khan et al., 1996) in pathological conditions. Interestingly, TGFβ seems to play an important role in mediating tendon repair (Chen et al., 2004) although the exact mechanism remains unclear. More recently, Kaji et al (Kaji et al., 2020) reported that TGFβ signaling is required in neonatal tenocytes for their recruitment to the site of tendon transection, and may play a role in promoting neonatal tendon regeneration.

Notably, this study identified a role for TGFβ signaling for recruitment of the resident tenocytes into the wound site and suggested a possible additional cell population involved in this process. In the present study we provide evidence for a distinctly different role for TGFβ signaling in tendon pathology, i.e. recruitment of a separate population of stem/progenitor cells from a distant niche into the degenerating tendon. Interestingly, these results highlight repeated involvement of TGFβ signaling in distinct cellular events in tendon biology.

Our results also indicate a cell autonomous requirement of TGFß signaling for the recruitment. In the Double Cre experiment with *RosaCreERT2*, the *Tgfbr2* receptor was eliminated from all cells. The failure of cell recruitment in this scenario could therefore be the result of a role for TGFβ signaling in the recruiting tendon, the environment surrounding the tendon or in the recruited cells themselves. However, as shown in Figure 8D, individual cells that did not lose receptor expression were recruited in this scenario, suggesting that the disruption was not in the tendon or tendon environment, since that would affect all cell recruitment, but rather a direct effect on the stem/progenitor cells that lost *Tgfbr2* expression. Future studies will focus on the mechanism of action at the cellular level of TGFβ signaling by transcriptome analysis of the recruited cells.

Lastly, TGFβ signaling is often associated with collagen matrix production (Klein et al., 2002; Leask and Abraham, 2004). However, apparent collagen disorganization was not observed in *Tgfbr2^ScxCre^* mutant tendons at the onset of the cellular phenotype (Tan et al., 2020). Instead, collagen disorganization and epitenon deterioration were noted only in pups older than one-week. The degenerative changes might thus imply a secondary consequence of the cellular changes in these mutants and/or of their movement difficulties. Regardless of the underlying causes, degenerative change has been implicated as a feature of tendinopathy (Kannus and Jozsa, 1991; Jarvinen et al., 1997; Longo et al., 2018). Much of what we learn about tendon healing comes from studies on acute tendon injury (Dyment et al., 2013; Wang et al., 2017; Kaji et al., 2020), and their significance to healing of tendinopathic tendons remains questionable. Moreover, it is difficult to obtain early tendinopathic human tissues because the conditions are often initially asymptomatic. The degenerative phenotype in *Tgfbr2^ScxCre^* mutant thus warrants further investigation and may provide a useful animal model for analysis of early degenerative changes in tendons.

## Materials and Methods

### Mice

Floxed TGFβ type II receptor (*Tgfbr2^f/f^*) mice (Chytil et al., 2002) were crossed with mice carrying the tendon deletor *Scleraxis-Cre* recombinase (*ScxCre*) (Blitz et al., 2013) to disrupt TGFβ signalling in tenocytes. The generation of *Rosa26-CreERT2 (RosaTCreERT2*) (Hameyer et al., 2007) and *Sox9CreERT2* (Soeda et al., 2010) mice have been described previously. All mice in this study also carried a transgenic tendon reporter *ScxGFP* (Pryce et al., 2007), and Cre reporters *Ai14 Rosa26-tdTomato (RosaT*) (Madisen et al., 2010) or *mTmG* (Muzumdar et al., 2007) for lineage tracing of *Scx*-expressing cells. All animal procedures were approved by the by the Animal Care and Use Committee at Oregon Health & Science University (OHSU). For embryo harvest, timed mating was set up in the afternoon, and identification of a mucosal plug on the next morning was considered 0.5 days of gestation (E0.5). Embryonic day 14.5 to postnatal day 13 (E14.5-P13) limb tendons were used for analysis. The mice genotype was determined by PCR analysis of DNA extracted from tail snip using a lysis reagent (Viagen Biotech, Cat 102-T) and proteinase K digestion (55°C, overnight).

### Transmission electron microscopy (TEM)

Mouse forelimbs were skinned, fixed in 1.5% glutaraldehyde/1.5% formaldehyde and decalcified in 0.2M EDTA. TEM was then performed as previously described (Tan et al., 2020). The acquired images were stitched using ImageJ software (https://imagej.nih.gov/ij/) (Preibisch et al., 2009).

### *In situ* hybridization and immunohistochemistry staining

Dissected forelimbs were fixed, decalcified and incubated with a 5-30% sucrose/PBS gradient. The tissues were then embedded in OCT (Tissue-Tek, Inc) and sectioned at 10-12 μm using a Microm HM550 cryostat (Thermo Scientific, Waltham, MA). In situ hybridization and immunohistochemistry staining were performed as previously described (Tan et al., 2020). For all qualitative studies, sections from two to four pups were examined to ensure reproducibility of results.

### Double Cre experiment to investigate the role of TGFβ signaling in cell recruitment

*CreERT2* recombinase was incorporated into the background of *Tgfbr2^f/-^;ScxCre* mutant to enable induction of either ubiquitous (*RosaCreERT2*) or restricted (*Sox9CreERT2) Tgfbr2* deletion upon tamoxifen administration. The pups were administered orally with tamoxifen (50 mg/ml in autoclaved corn oil; 15 μl per pup) at P1 and P2 and harvested at P7, a stage at which only recruited cells are positive for *ScxGFP* in the mutant tendon.

### Quantification of recruited cells in transverse sections of the communis tendon

For the analysis of the effect of TGFβ signaling ablation on cell recruitment, the number of recruited cells were counted in extensor digitorium communis tendons at the zeugopod level of both P7 *Tgfbr2^ScxCre^* and Double Cre (*ScxCre;RosaCreERT* and *ScxCre;Sox9CreERT*) mutant pups. Forelimb transverse sections were stained DAPI to highlight the nuclei for identifying the individual cell. Recruited cells were identified by positive *ScxGFP* signal at this stage and their number was counted in 6-15 sections per pup from 3-5 pups.

### Statistical analysis

Student’s *t*-tests were performed to determine the statistical significance of differences between groups (n ≥ 3). Unless stated otherwise, all graphs are presented as mean ± standard deviation (SD). A value of *p*<0.05 is regarded as statistically significant.

## Acknowledgement

The authors thank Dr Elazar Zelzer (Department of Molecular Genetics, Weizmann Institute of Science, Israel) for critical reading of the manuscript. We are grateful to staff from FACS Core, OHSU particularly Dr Miranda Gilchrist for their excellent technical assistance. This work was funded by NIH (R01AR055973) and Shriners Hospital for Children (SHC 5410-POR-14). G.K.T was supported by Research Fellowship from Shriners Hospital for Children.

---

**Supplementary Figure 1.**
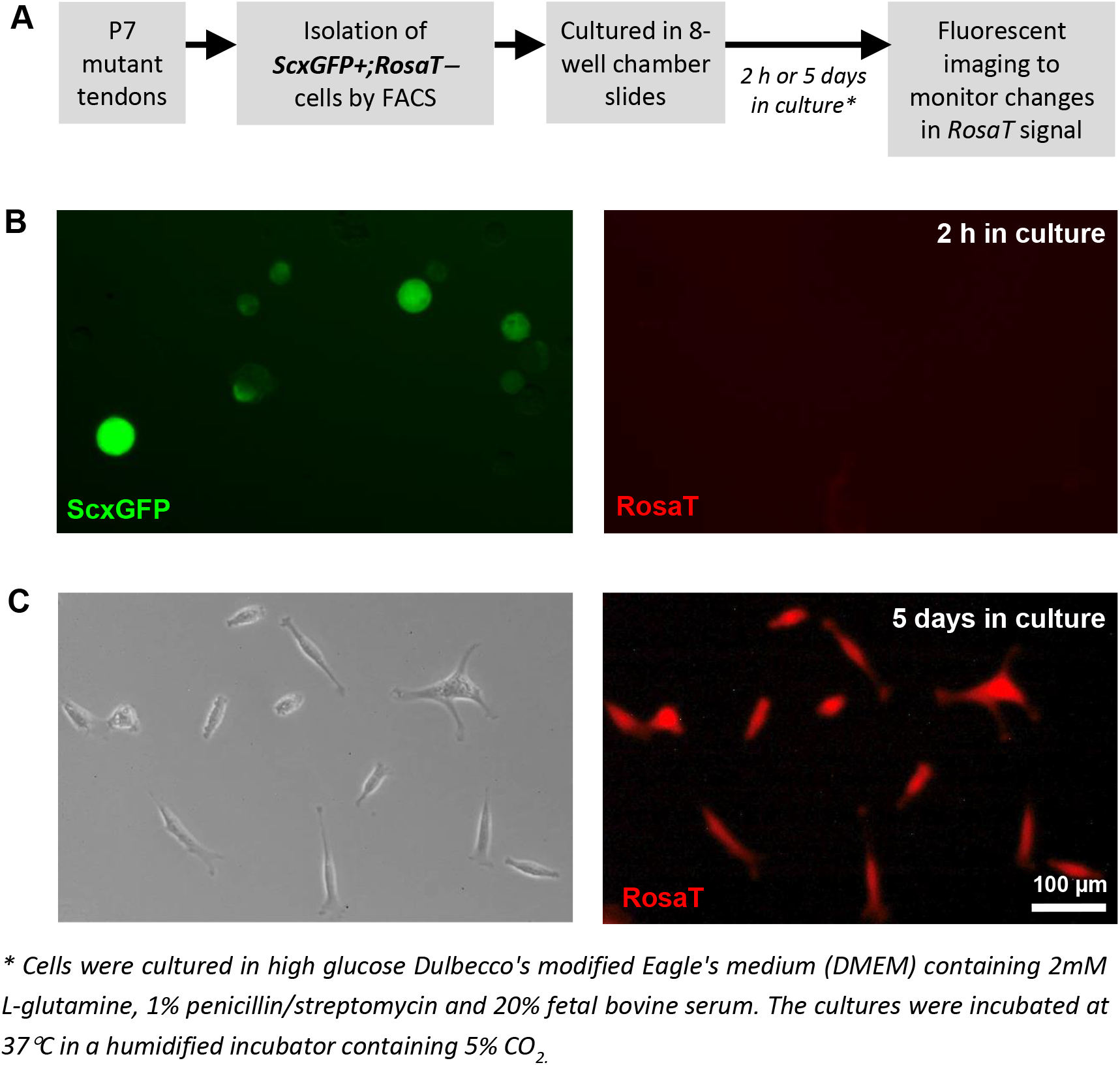
Newly recruited *ScxGFP+;RosaT−* cells subsequently induce expression of the Cre reporter. (A) Experimental outline: Cells were enzymatically-released from P7 *Tgfbr2^ScxCre^* mutant tendons. *ScxGFP+;RosaT−*cells were then sorted by FACS and cultured in 8-well chamber slides. Fluorescence imaging was carried out at designated time-points to monitor changes in *RosaT* signal. (B) Fluorescent images of the sorted cells after 2 h in culture. Note that most cells were still floating in medium. As determined by the FACS gating, only *ScxGFP*-positive and *RosaT*-negative cells were isolated. Images not to scale. (C) Brightfield and fluorescent images of the sorted cells after 5 days of culture, in which all of the cells have now expressed *RosaT* Cre reporter. These findings support the hypothesis that, upon induction of the *Scx* enhancer, the *ScxGFP* signal will be detected first and the *RosaT* signal will follow with some delay.

**Supplementary Figure 2.**
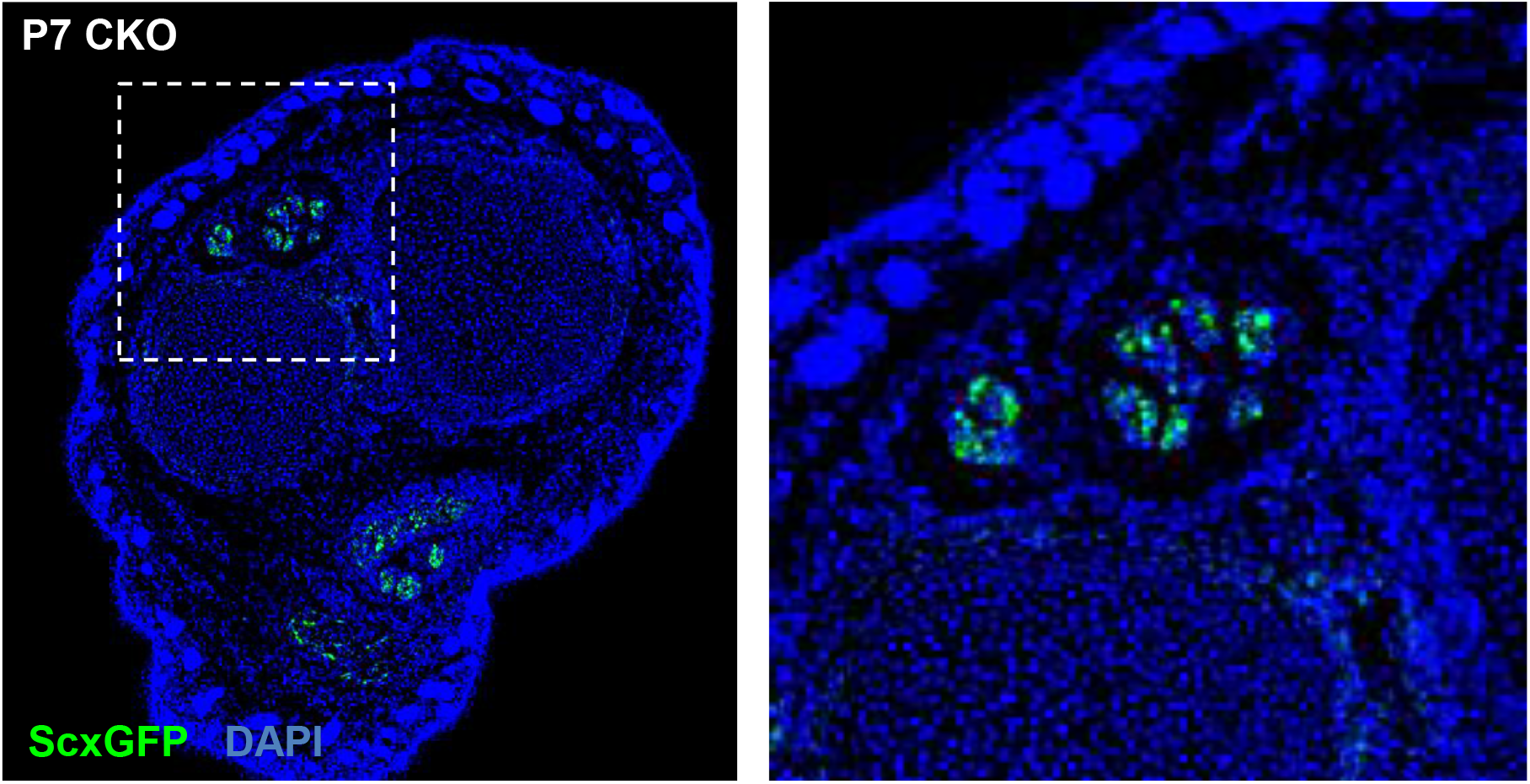
The recruited cells turned on tendon gene expression only upon arrival to or entering the tendon. Histological examination of transverse sections of *Tgfbr2^ScxCre^* mutant forelimbs stained with DAPI to visualize cell distribution in the section and *ScxGFP* to identify the recruited cells. Shown here is a representative histological section from P7 mutants. Recruited cells expressing the tendon reporter *ScxGFP* were not detected away from the tendons but only within or adjacent to mutant tendons, suggesting that irrespective of their origin the cells turned on tendon gene expression only upon arrival to or entering the tissue. Boxed region is shown enlarged in the right panel. Images are not to scale.

## References

Beekman, C., Nichane, M., De Clercq, S., Maetens, M., Floss, T., Wurst, W., Bellefroid, E. and Marine, J. C. (2006) ‘Evolutionarily conserved role of nucleostemin: controlling proliferation of stem/progenitor cells during early vertebrate development’, Molecular and cellular biology 26(24): 9291–301.

Bi, Y., Ehirchiou, D., Kilts, T. M., Inkson, C. A., Embree, M. C., Sonoyama, W., Li, L., Leet, A. I., Seo, B. M., Zhang, L. et al. (2007) ‘Identification of tendon stem/progenitor cells and the role of the extracellular matrix in their niche’, Nature medicine 13(10): 1219–27.

Biro, T. and Bihari-Varga, M. (1974) ‘[Thermoanalytic studies in connective tissue research. II. Changes in the structure of cartilage and tendon tissues under pathologic conditions]’, Magyar traumatologia, orthopaedia es helyreallito sebeszet 17(4): 282–6.

Blitz, E., Sharir, A., Akiyama, H. and Zelzer, E. (2013) ‘Tendon-bone attachment unit is formed modularly by a distinct pool of Scx- and Sox9-positive progenitors’, Development 140(13): 2680–90.

Boileau, P., Brassart, N., Watkinson, D. J., Carles, M., Hatzidakis, A. M. and Krishnan, S. G. (2005) ‘Arthroscopic repair of full-thickness tears of the supraspinatus: does the tendon really heal?’, The Journal of bone and joint surgery. American volume 87(6): 1229–40.

Cathery, W., Faulkner, A., Maselli, D. and Madeddu, P. (2018) ‘Concise Review: The Regenerative Journey of Pericytes Toward Clinical Translation’, Stem cells 36(9): 1295–1310.

Chen, Y. J., Wang, C. J., Yang, K. D., Kuo, Y. R., Huang, H. C., Huang, Y. T., Sun, Y. C. and Wang, F. S. (2004) ‘Extracorporeal shock waves promote healing of collagenase-induced Achilles tendinitis and increase TGF-beta1 and IGF-I expression’, Journal of orthopaedic research : official publication of the Orthopaedic Research Society 22(4): 854–61.

Chytil, A., Magnuson, M. A., Wright, C. V. and Moses, H. L. (2002) ‘Conditional inactivation of the TGF-beta type II receptor using Cre:Lox’, Genesis 32(2): 73–5.

Cook, J. L., Rio, E., Purdam, C. R. and Docking, S. I. (2016) ‘Revisiting the continuum model of tendon pathology: what is its merit in clinical practice and research?’, Br J Sports Med 50(19): 1187–91.

Dahlgren, L. A., Mohammed, H. O. and Nixon, A. J. (2005) ‘Temporal expression of growth factors and matrix molecules in healing tendon lesions’, Journal of orthopaedic research : official publication of the Orthopaedic Research Society 23(1): 84–92.

Dyment, N. A., Hagiwara, Y., Matthews, B. G., Li, Y., Kalajzic, I. and Rowe, D. W. (2014) ‘Lineage tracing of resident tendon progenitor cells during growth and natural healing’, PLoS One 9(4): e96113.

Dyment, N. A., Liu, C. F., Kazemi, N., Aschbacher-Smith, L. E., Kenter, K., Breidenbach, A. P., Shearn, J. T., Wylie, C., Rowe, D. W. and Butler, D. L. (2013) ‘The paratenon contributes to scleraxis-expressing cells during patellar tendon healing’, PLoS One 8(3): e59944.

Favata, M., Beredjiklian, P. K., Zgonis, M. H., Beason, D. P., Crombleholme, T. M., Jawad, A. F. and Soslowsky, L. J. (2006) ‘Regenerative properties of fetal sheep tendon are not adversely affected by transplantation into an adult environment’, Journal of orthopaedic research : official publication of the Orthopaedic Research Society24(11): 2124–32.

Fenwick, S. A., Curry, V., Harrall, R. L., Hazleman, B. L., Hackney, R. and Riley, G. P. (2001) ‘Expression of transforming growth factor-beta isoforms and their receptors in chronic tendinosis’, Journal of anatomy 199(Pt 3): 231–40.

Ferry, S. T., Dahners, L. E., Afshari, H. M. and Weinhold, P. S. (2007) ‘The effects of common antiinflammatory drugs on the healing rat patellar tendon’, The American journal of sports medicine35(8): 1326–33.

Formeister, E. J., Sionas, A. L., Lorance, D. K., Barkley, C. L., Lee, G. H. and Magness, S. T. (2009) ‘Distinct SOX9 levels differentially mark stem/progenitor populations and enteroendocrine cells of the small intestine epithelium’, American journal of physiology. Gastrointestinal and liver physiology 296(5): G1108–18.

Forslund, C. and Aspenberg, P. (2003) ‘Improved healing of transected rabbit Achilles tendon after a single injection of cartilage-derived morphogenetic protein-2’, The American journal of sports medicine 31(4): 555–9.

Franitza, S., Kollet, O., Brill, A., Vaday, G. G., Petit, I., Lapidot, T., Alon, R. and Lider, O. (2002) ‘TGF-beta1 enhances SDF-1alpha-induced chemotaxis and homing of naive T cells by up-regulating CXCR4 expression and downstream cytoskeletal effector molecules’, European journal of immunology 32(1): 193–202.

Furuyama, K., Kawaguchi, Y., Akiyama, H., Horiguchi, M., Kodama, S., Kuhara, T., Hosokawa, S., Elbahrawy, A., Soeda, T., Koizumi, M. et al. (2011) ‘Continuous cell supply from a Sox9-expressing progenitor zone in adult liver, exocrine pancreas and intestine’, Nature genetics 43(1): 34–41.

Gerber, C., Fuchs, B. and Hodler, J. (2000) ‘The results of repair of massive tears of the rotator cuff’, The Journal of bone and joint surgery. American volume 82(4): 505–15.

Hameyer, D., Loonstra, A., Eshkind, L., Schmitt, S., Antunes, C., Groen, A., Bindels, E., Jonkers, J., Krimpenfort, P., Meuwissen, R. et al. (2007) ‘Toxicity of ligand-dependent Cre recombinases and generation of a conditional Cre deleter mouse allowing mosaic recombination in peripheral tissues’, Physiological genomics 31(1): 32–41.

Harrison, R. K., Mudera, V., Grobbelaar, A. O., Jones, M. E. and McGrouther, D. A. (2003) ‘Synovial sheath cell migratory response to flexor tendon injury: an experimental study in rats’, The Journal of hand surgery 28(6): 987–93.

Harvey, T., Flamenco, S. and Fan, C. M. (2019) ‘A Tppp3(+)Pdgfra(+) tendon stem cell population contributes to regeneration and reveals a shared role for PDGF signalling in regeneration and fibrosis’, Nature cell biology 21(12): 1490–1503.

Havis, E., Bonnin, M. A., Olivera-Martinez, I., Nazaret, N., Ruggiu, M., Weibel, J., Durand, C., Guerquin, M. J., Bonod-Bidaud, C., Ruggiero, F. et al. (2014) ‘Transcriptomic analysis of mouse limb tendon cells during development’, Development 141(19): 3683–96.

He, X., Bougioukli, S., Ortega, B., Arevalo, E., Lieberman, J. R. and McMahon, A. P. (2017) ‘Sox9 positive periosteal cells in fracture repair of the adult mammalian long bone’, Bone 103: 12–19.

Hernigou, P., Flouzat Lachaniette, C. H., Delambre, J., Zilber, S., Duffiet, P., Chevallier, N. and Rouard, H. (2014) ‘Biologic augmentation of rotator cuff repair with mesenchymal stem cells during arthroscopy improves healing and prevents further tears: a case-controlled study’, Int Orthop 38(9): 1811–8.

Howell, K., Chien, C., Bell, R., Laudier, D., Tufa, S. F., Keene, D. R., Andarawis-Puri, N. and Huang, A. H. (2017) ‘Novel Model of Tendon Regeneration Reveals Distinct Cell Mechanisms Underlying Regenerative and Fibrotic Tendon Healing’, Scientific reports 7: 45238.

Huang, A. H., Watson, S. S., Wang, L., Baker, B. M., Akiyama, H., Brigande, J. V. and Schweitzer, R. (2019) ‘Requirement for scleraxis in the recruitment of mesenchymal progenitors during embryonic tendon elongation’, Development 146(20).

Jarvinen, M., Jozsa, L., Kannus, P., Jarvinen, T. L., Kvist, M. and Leadbetter, W. (1997) ‘Histopathological findings in chronic tendon disorders’, Scandinavian journal of medicine & science in sports 7(2): 86–95.

Jones, M. E., Mudera, V., Brown, R. A., Cambrey, A. D., Grobbelaar, A. O. and McGrouther, D. A. (2003) ‘The early surface cell response to flexor tendon injury’, The Journal of hand surgery 28(2): 221–30.

Jujo, K., Hamada, H., Iwakura, A., Thorne, T., Sekiguchi, H., Clarke, T., Ito, A., Misener, S., Tanaka, T., Klyachko, E. et al. (2010) ‘CXCR4 blockade augments bone marrow progenitor cell recruitment to the neovasculature and reduces mortality after myocardial infarction’, Proceedings of the National Academy of Sciences of the United States of America 107(24): 11008–13.

Kaji, D. A., Howell, K. L., Balic, Z., Hubmacher, D. and Huang, A. H. (2020) ‘Tgfbeta signaling is required for tenocyte recruitment and functional neonatal tendon regeneration’, Elife 9.

Kan, C., Chen, L., Hu, Y., Ding, N., Li, Y., McGuire, T. L., Lu, H., Kessler, J. A. and Kan, L. (2018) ‘Gli1-labeled adult mesenchymal stem/progenitor cells and hedgehog signaling contribute to endochondral heterotopic ossification’, Bone 109: 71–79.

Kannus, P. (2000) ‘Structure of the tendon connective tissue’, Scandinavian journal of medicine & science in sports 10(6): 312–20.

Kannus, P. and Jozsa, L. (1991) ‘Histopathological changes preceding spontaneous rupture of a tendon. A controlled study of 891 patients’, The Journal of bone and joint surgery. American volume 73(10): 1507–25.

Khan, U., Edwards, J. C. and McGrouther, D. A. (1996) ‘Patterns of cellular activation after tendon injury’, J Hand Surg Br 21(6): 813–20.

Klein, M. B., Yalamanchi, N., Pham, H., Longaker, M. T. and Chang, J. (2002) ‘Flexor tendon healing in vitro: effects of TGF-beta on tendon cell collagen production’, The Journal of hand surgery 27(4): 615–20.

Kragsnaes, M. S., Fredberg, U., Stribolt, K., Kjaer, S. G., Bendix, K. and Ellingsen, T. (2014) ‘Stereological quantification of immune-competent cells in baseline biopsy specimens from achilles tendons: results from patients with chronic tendinopathy followed for more than 4 years’, The American journal of sports medicine 42(10): 2435–45.

Leask, A. and Abraham, D. J. (2004) ‘TGF-beta signaling and the fibrotic response’, FASEB journal: official publication of the Federation of American Societies for Experimental Biology 18(7): 816–27.

Longo, U. G., Franceschi, F., Ruzzini, L., Rabitti, C., Morini, S., Maffulli, N. and Denaro, V. (2008) ‘Histopathology of the supraspinatus tendon in rotator cuff tears’, The American journal of sports medicine 36(3): 533–8.

Longo, U. G., Ronga, M. and Maffulli, N. (2018) ‘Achilles Tendinopathy’, Sports Med Arthrosc Rev26(1): 16–30.

Lui, P. P. (2015) ‘Markers for the identification of tendon-derived stem cells in vitro and tendon stem cells in situ - update and future development’, Stem cell research & therapy 6: 106.

Madisen, L., Zwingman, T. A., Sunkin, S. M., Oh, S. W., Zariwala, H. A., Gu, H., Ng, L. L., Palmiter, R. D., Hawrylycz, M. J., Jones, A. R. et al. (2010) ‘A robust and high-throughput Cre reporting and characterization system for the whole mouse brain’, Nature neuroscience 13(1): 133–40.

Marecic, O., Tevlin, R., McArdle, A., Seo, E. Y., Wearda, T., Duldulao, C., Walmsley, G. G., Nguyen, A., Weissman, I. L., Chan, C. K. et al. (2015) ‘Identification and characterization of an injury-induced skeletal progenitor’, Proceedings of the National Academy of Sciences of the United States of America 112(32): 9920–5.

Mathews, V., Hanson, P. T., Ford, E., Fujita, J., Polonsky, K. S. and Graubert, T. A. (2004) ‘Recruitment of bone marrow-derived endothelial cells to sites of pancreatic beta-cell injury’,Diabetes 53(1): 91–8.

Mendias, C. L., Gumucio, J. P., Bakhurin, K. I., Lynch, E. B. and Brooks, S. V. (2012) ‘Physiological loading of tendons induces scleraxis expression in epitenon fibroblasts’, J Orthop Res 30(4): 606–12.

Mienaltowski, M. J., Adams, S. M. and Birk, D. E. (2013) ‘Regional differences in stem cell/progenitor cell populations from the mouse achilles tendon’, Tissue engineering. Part A 19(1-2): 199–210.

Millar, N. L., Hueber, A. J., Reilly, J. H., Xu, Y., Fazzi, U. G., Murrell, G. A. and McInnes, I. B. (2010) ‘Inflammation is present in early human tendinopathy’, The American journal of sports medicine38(10): 2085–91.

Muzumdar, M. D., Tasic, B., Miyamichi, K., Li, L. and Luo, L. (2007) ‘A global double-fluorescent Cre reporter mouse’, Genesis 45(9): 593–605.

Nomura, J., Maruyama, M., Katano, M., Kato, H., Zhang, J., Masui, S., Mizuno, Y., Okazaki, Y., Nishimoto, M. and Okuda, A. (2009) ‘Differential requirement for nucleostemin in embryonic stem cell and neural stem cell viability’, Stem cells 27(5): 1066–76.

Nourissat, G., Diop, A., Maurel, N., Salvat, C., Dumont, S., Pigenet, A., Gosset, M., Houard, X. and Berenbaum, F. (2010) ‘Mesenchymal stem cell therapy regenerates the native bone-tendon junction after surgical repair in a degenerative rat model’, PLoS One 5(8): e12248.

Oh, J. H., Chung, S. W., Kim, S. H., Chung, J. Y. and Kim, J. Y. (2014) ‘2013 Neer Award: Effect of the adipose-derived stem cell for the improvement of fatty degeneration and rotator cuff healing in rabbit model’, Journal of shoulder and elbow surgery 23(4): 445–55.

Pineault, K. M., Song, J. Y., Kozloff, K. M., Lucas, D. and Wellik, D. M. (2019) ‘Hox11 expressing regional skeletal stem cells are progenitors for osteoblasts, chondrocytes and adipocytes throughout life’, Nature communications 10(1): 3168.

Poche, R. A., Furuta, Y., Chaboissier, M. C., Schedl, A. and Behringer, R. R. (2008) ‘Sox9 is expressed in mouse multipotent retinal progenitor cells and functions in Muller glial cell development’, The Journal of comparative neurology 510(3): 237–50.

Preibisch, S., Saalfeld, S. and Tomancak, P. (2009) ‘Globally optimal stitching of tiled 3D microscopic image acquisitions’, Bioinformatics 25(11): 1463–5.

Pryce, B. A., Brent, A. E., Murchison, N. D., Tabin, C. J. and Schweitzer, R. (2007) ‘Generation of transgenic tendon reporters, ScxGFP and ScxAP, using regulatory elements of the scleraxis gene’, Developmental dynamics : an official publication of the American Association of Anatomists 236(6): 1677–82.

Rennert, R. C., Sorkin, M., Garg, R. K. and Gurtner, G. C. (2012) ‘Stem cell recruitment after injury: lessons for regenerative medicine’, Regenerative medicine 7(6): 833–50.

Rinkevich, Y., Lindau, P., Ueno, H., Longaker, M. T. and Weissman, I. L. (2011) ‘Germ-layer and lineage-restricted stem/progenitors regenerate the mouse digit tip’, Nature 476(7361): 409–13.

Runesson, E., Ackermann, P., Karlsson, J. and Eriksson, B. I. (2015) ‘Nucleostemin- and Oct 3/4-positive stem/progenitor cells exhibit disparate anatomical and temporal expression during rat Achilles tendon healing’, BMC Musculoskelet Disord 16: 212.

Sakabe, T., Sakai, K., Maeda, T., Sunaga, A., Furuta, N., Schweitzer, R., Sasaki, T. and Sakai, T. (2018) ‘Transcription factor scleraxis vitally contributes to progenitor lineage direction in wound healing of adult tendon in mice’, The Journal of biological chemistry 293(16): 5766–5780.

Scott, C. E., Wynn, S. L., Sesay, A., Cruz, C., Cheung, M., Gomez Gaviro, M. V., Booth, S., Gao, B., Cheah, K. S., Lovell-Badge, R. et al. (2010) ‘SOX9 induces and maintains neural stem cells’, Nature neuroscience 13(10): 1181–9.

Sharma, P. and Maffulli, N. (2005) ‘Tendon injury and tendinopathy: healing and repair’, The Journal of bone and joint surgery. American volume 87(1): 187–202.

Soeda, T., Deng, J. M., de Crombrugghe, B., Behringer, R. R., Nakamura, T. and Akiyama, H. (2010) ‘Sox9-expressing precursors are the cellular origin of the cruciate ligament of the knee joint and the limb tendons’, Genesis 48(11): 635–44.

Tan, G. K., Pryce, B. A., Stabio, A., Brigande, J. V., Wang, C., Xia, Z., Tufa, S. F., Keene, D. R. and Schweitzer, R. (2020) ‘Tgfbeta signaling is critical for maintenance of the tendon cell fate’, Elife 9.

Tan, Q., Lui, P. P. and Lee, Y. W. (2013) ‘In vivo identity of tendon stem cells and the roles of stem cells in tendon healing’, Stem Cells Dev 22(23): 3128–40.

Tang, Y., Wu, X., Lei, W., Pang, L., Wan, C., Shi, Z., Zhao, L., Nagy, T. R., Peng, X., Hu, J. et al. (2009) ‘TGF-beta1-induced migration of bone mesenchymal stem cells couples bone resorption with formation’, Nature medicine 15(7): 757–65.

Wan, M., Li, C., Zhen, G., Jiao, K., He, W., Jia, X., Wang, W., Shi, C., Xing, Q., Chen, Y. F. et al. (2012) ‘Injury-activated transforming growth factor beta controls mobilization of mesenchymal stem cells for tissue remodeling’, Stem cells 30(11): 2498–511.

Wang, Y., Zhang, X., Huang, H., Xia, Y., Yao, Y., Mak, A. F., Yung, P. S., Chan, K. M., Wang, L., Zhang, C. et al. (2017) ‘Osteocalcin expressing cells from tendon sheaths in mice contribute to tendon repair by activating Hedgehog signaling’, Elife 6.

Xynos, A., Corbella, P., Belmonte, N., Zini, R., Manfredini, R. and Ferrari, G. (2010) ‘Bone marrow-derived hematopoietic cells undergo myogenic differentiation following a Pax-7 independent pathway’, Stem cells 28(5): 965–73.

Yin, H., Price, F. and Rudnicki, M. A. (2013) ‘Satellite cells and the muscle stem cell niche’, Physiological reviews 93(1): 23–67.

Zhang, J. and Wang, J. H. (2010) ‘Characterization of differential properties of rabbit tendon stem cells and tenocytes’, BMC musculoskeletal disorders 11: 10.

